# Extracellular Matrix Regulates Neuronal Chloride Concentration via K^+^-Cl^−^-Cotransporter 2

**DOI:** 10.1101/2023.02.09.527837

**Authors:** Tushar Devanand Yelhekar, Tatiana Kuznetsova, Evgenya Malinina, Evgeni Ponimaskin, Alexander Dityatev, Michael Druzin, Staffan Johansson

## Abstract

The neuronal intracellular chloride concentration [Cl^−^]_i_ is critical for γ‐aminobutyric acid type A (GABA_A_) receptor‐mediated transmission. Degradation of the extracellular matrix (ECM) is associated with raised [Cl^−^]_I_ but neither the mechanisms underlying this effect nor the consequences for GABA‐mediated transmission are well understood. Hitherto it has been unclear how to reconcile the effect of the ECM on [Cl^−^]_i_ with the established role of cation‐chloride cotransporters in setting [Cl^−^]_I_. In the present work we clarify the role of the ECM in the control of neuronal [Cl^−^]_i_. By measuring [Cl^−^]_i_ in central neurons from male rats we show that the ECM affects basal [Cl^−^]_i_ as well as the rate of Cl^−^ extrusion after a high load. The mechanism is not via impermeant anions but through regulation of K^+^‐Cl^−^‐cotransporter 2 (KCC2). ECM degradation is accompanied by an N‐type Ca^2+^‐channel‐ and calpain‐dependent reduction in the amount of KCC2 protein, increased basal [Cl^−^]_i_, reduced Cl^−^ extrusion capacity as well as by reduced inhibitory, or even an excitatory, effect of intense GABA_A_‐ receptor mediated trans mission. This implies a previously unrecognized pathway for the control of neuronal [Cl^−^]_i_ and excitability by the ECM.

**Graphical abstract:** 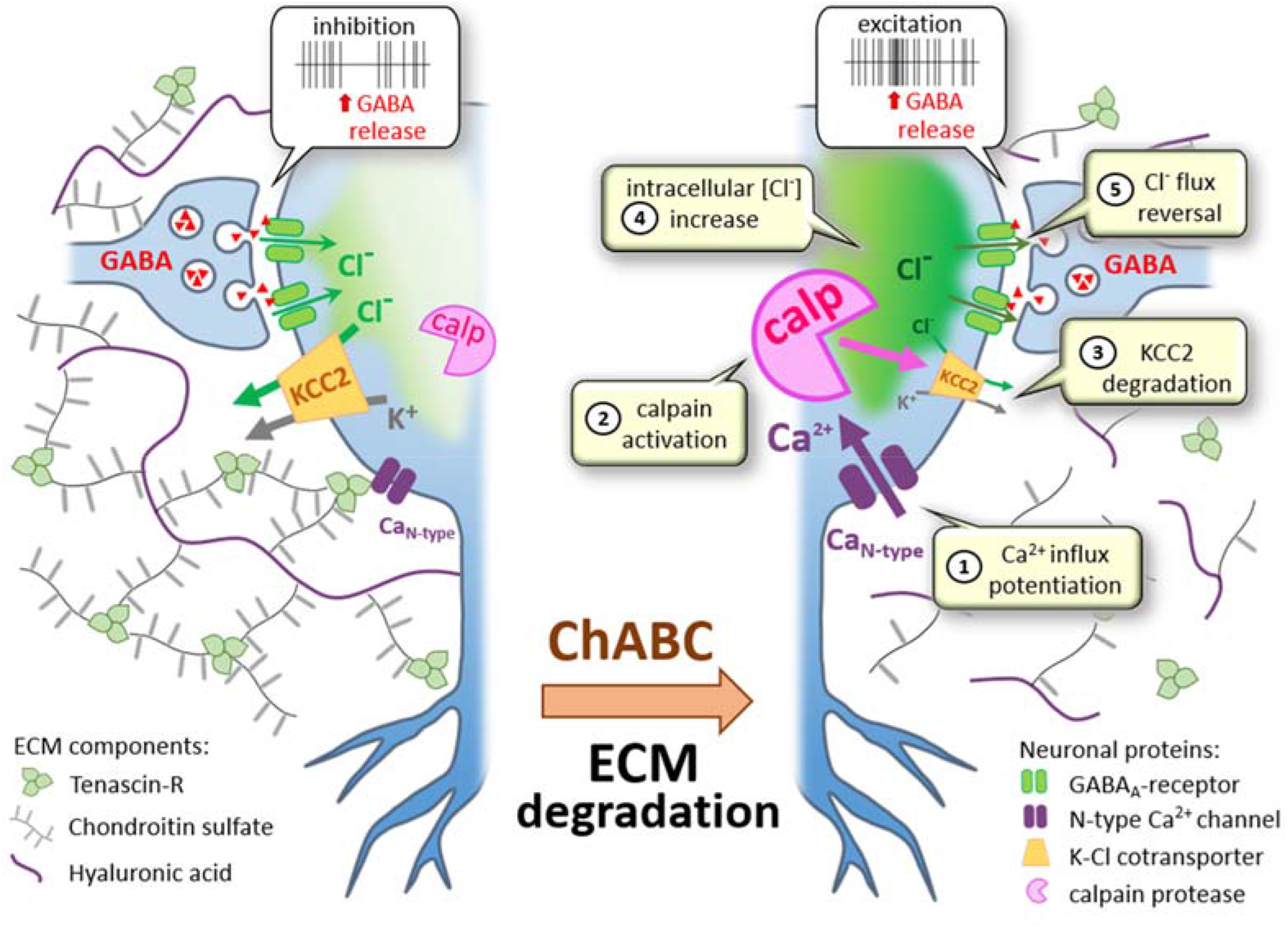

## Introduction

GABA_A_ receptor‐mediated neurotransmission is critical for normal brain function. The GABA_A_ receptor‐channel is permeable to Cl^−^ and thus the effect of activation depends critically on the intracellular Cl^−^ concentration ([Cl^−^]_i_). The inhibitory action of G_A_BA_A_ receptors on neuronal excitability that predominates in the adult brain is a consequence of a low [Cl^−^]_i_ (often 5 – 10 mM). Higher [Cl^−^]_i_ is found early in development (Ben‐Ari, 2002), in some adult neurons (Choi et al., 2008; DeFazio et al., 2002; Haam et al., 2012) and in pathological conditions such as epilepsy (Huberfeld et al., 2007) and chronic pain (Coull et al., 2003) and may be associated with an excitatory effect of GABA_A_‐receptor activation. The extracellular matrix (ECM), which plays important roles in synaptic plasticity and is proteolytically degraded during epileptogenesis (Soleman et al., 2013; Senkov et al., 2014), may also be important for the control of basal [Cl^−^]_i_. Thus, it has been shown that ECM degradation is associated with increased [Cl^−^]_i_ (Glykys et al., 2014a). The mechanism first suggested to explain this effect involved negative charges maintained by the ECM (Glykys et al., 2014a). However, the proposal questioned the previously ascribed role of the cation‐chloride cotransporters, Na^+^‐K^+^‐Cl^−^‐cotransporter 1 (NKCC1) and K^+^‐Cl^−^‐cotransporter 2 (KCC2) and has been questioned and remains controversial (Ben‐Ari, 2014; Cho, 2014; Delpire and Staley, 2014; Glykys et al., 2014b; Kaila et al., 2014; Luhmann et al., 2014; Voipio et al., 2014; Savtchenko et al., 2017). NKCC1 and KCC2 remain seen as the main controllers of [Cl^−^]_i_. While NKCC1, if present, transports Cl^−^ into the cell, KCC2 transports Cl^−^ mainly out of the cell, but may contribute to inward Cl^−^ transport depending on the driving force set by the concentration of transported ions. KCC2 has been thought necessary to maintain the low Cl^−^]_i_ required for the common type of hyperpolarizing responses, including inhibitory post‐synaptic potentials, to GABA and glycine observed in mature central neurons (Kaila et al., 2014). The question how the ECM affects [Cl^−^]_I_ and determines the inhibitory/excitatory effect of GABA_A_‐receptor activation has remained unanswered.

In the present work, we analyze the mechanism by which the ECM influences [Cl^−^]_i_ and how it affects GABA_A_‐receptor mediated transmission and neuronal excitability. By estimating [Cl^−^]_i_ directly from electrophysiological recordings of Cl^−^ currents, we show that the integrity of the ECM is important for the basal [Cl^−^]_i_ and also that the ECM strongly influences the rate of Cl^−^ extrusion after a high Cl^−^ load. Our results, however, are not compatible with a mechanism involving a strong effect of negative charges of the ECM (Glykys et al., 2014a). Rather, our combined electrophysiological and biochemical results show that degradation of the ECM alters the amount of KCC2 via Ca^2+^ influx through N‐type channels and calpain activation, thus regulating the rate of KCC2‐mediated Cl^−^ extrusion. Thereby the ECM exerts an influence on the postsynaptic effects mediated by GABA_A_‐receptor activation and preserves an inhibitory action in the presence of intense neuronal activity.

## Materials and Methods

### Ethical approval

Ethical approval of the procedures described was given either by the regional ethics committee for animal research (“Umeå djurförsöksetiska nämnd”, approvals no. A18‐11 and A9‐14) or by the Hannover Medical School Institutional Animal Care and Research Advisory Committee, with all experiments permitted by the local government (#33.14 42502‐04‐12/0753).

### Brain slice preparation

Male Sprague‐Dawley rats, aged 3 – 5 weeks, were used for the experiments. The procedures to prepare 200 – 300 µm thick coronal brain slices containing the preoptic area have been described elsewhere (Karlsson et al., 2011, modified after Malinina et al., 2005).

### Chondroitinase ABC treatment

In some of the brain slices used, degradation of ECM was induced by chondroitinase ABC treatment (ChABC) (0.2 U ml^−1^; AMS Biotechnology) for at least 2 h at about 36 °C. ChABC was dissolved in standard extracellular solution containing 0.1 % bovine serum albumin (BSA; Sigma‐Aldrich). This protocol for ChABC treatment has proved functional for slice preparations of brain tissue (Bukalo et al., 2001; Hayani et al., 2018). The lower concentration of ChABC compared to that used e. g. by Glykys et al. (2014a) is likely of advantage for minimizing possible non‐specific side effects. Histochemical verification of ECM degradation was performed as described below. Control slices were exposed to similar treatment without ChABC. In some experiments, additional test substances, as specified below, were present during ChABC exposure.

### Histochemistry

Brain slices were divided in two cerebral hemispheres, for control and ChABC treatment (see above), respectively. The most lateral part of cerebral cortex in ChABC‐treated hemisections were cut off for identification. After treatment, slices were fixed overnight in 4 % formaldehyde in phosphate‐buffered saline (PBS; Sigma‐Aldrich). Following wash in PBS, slices were incubated in PBS‐based cryoprotectant solutions: 2×10 min in 10 % sucrose (w/v), 2×20 min in 20 % sucrose with 6 % glycerol, then 2×30 min in 30 % sucrose and 12 % glycerol. Subsequently, the sections were carefully placed in envelopes of aluminium foil for permeabilization of cell membranes by rapid freeze–thaw cycles (Hughes et al., 2000). The slices were frozen by placement close above liquid nitrogen for 45 s and thawed until all resulting condensation on the aluminium had disappeared. This freeze–thaw cycle was repeated three times for each slice. After permeabilization, slices were rinsed three times in 0.25 % BSA in PBS for 20 min each and then incubated in PBS containing 5.0 % BSA and 0.2 % Triton X‐100 (Sigma‐Aldrich) for 1 h. Subsequently, slices were incubated with biotinylated *Wisteria floribunda* agglutinin (WFA; 1:400; Sigma‐Aldrich), diluted in PBS containing 2.5 % BSA. For negative control slices, WFA was omitted. After incubation for 3 days at 4°C at a shaking platform, slices were rinsed three to five times in PBS and exposed to streptavidin‐Alexa 488 (1:300; Thermo Fisher) overnight at 4°C. After three washes, slices were embedded in Vectashield mounting medium with 4’,6‐diamidino‐2‐phenylindole (DAPI; Vector Laboratories). Paired hemisections from ChABC‐treated and control groups were incubated together in one well, both in primary and in secondary reagent mixtures, and mounted together under one cover slip.

For an overall view of slices and to estimate the effect of ChABC on WFA reactivity (Howell et al., 2015; Morita et al., 2010), we used a conventional upright fluorescence microscope (Nikon Eclipse 80i) with a DS‐U2 digital camera. Images were acquired with a 1.6× air objective using FITC (GFP) filter (Ext 482/25, EM 530/40) for visualization of chondroitin sulphate proteoglycans, stained by WFA. The exposure time for each filter was constant to enable comparison of signals in control and experimental groups. The background fluorescence (average intensity in the negative control slice) was subtracted from each image. To obtain images of full hemisections, separate images of neighboring regions were acquired with overlap and then stitched using MosaicJ plugin of the Fiji software.

To view perineuronal nets (PNNs) in control sections (as a positive control for WFA staining; Härtig et al., 1992), we used an inverted confocal microscope (Nikon A1R). High‐magnification images were taken with a differential interference contrast 20× air (NA 0.75) objective. To visualize nuclei, DAPI was excited by 405 nm laser and to visualize PNNs, Alexa488 was excited by 488 nm laser. Selected areas were captured by a 20× objective in a single focal plane. Acquisition as well as processing of all images was made in the same way. Image processing was carried out using Fiji software.

### Recording solutions for electrophysiology

Unless stated otherwise, all solution chemicals were obtained from Sigma‐Aldrich. The standard extracellular solution contained (in mM): NaCl 137, KCl 5.0, CaCl_2_ 1.0, MgCl_2_ 1.2, HEPES 10, glucose 10, pH 7.4 (adjusted with NaOH). An alternative solution where some Na‐acetate was substituted for NaCl, to obtain a Cl^−^ concentration ([Cl^−^]_o_bulk_) of 50 mM, was used in a subset of experiments aimed at estimating local [Cl^−^]_o_ contributing to the driving force for Cl^−^.

For gramicidin‐perforated‐patch recordings, used to estimate [Cl^−^]_i_, the standard pipette‐filling solution contained 300 µg ml^−1^ gramicidin (Sigma‐Aldrich) and (in mM): K‐acetate 140, NaCl 3.0, MgCl_2_ 1.2, HEPES 10, EGTA 1.0, pH 7.2 (adjusted with KOH). An alternative high‐[Cl^−^] solution contained 300 µg ml^−1^ gramicidin and (in mM): KCl 140, NaCl, 3.0, MgCl_2_ 1.2, EGTA 1.0, HEPES 10, pH 7.2 (adjusted with KOH). The choice of pipette‐filling solution was made as to facilitate detection of patch rupture, with high‐[Cl^−^] solutions being used preferentially in experiments when [Cl^−^]_i_ was expected to be mainly at a low level and low‐[Cl^−^] solutions preferentially where [Cl^−^]_i_ was expected to be raised for prolonged periods. (In practice, however, this precaution was unnecessary, since whenever possible rupture was suspected, this could easily be verified or disproven by controlling the possibility to “set [Cl^−^]_i_” at high or low level by combined voltage control and GABA application; cf the “loading procedure” described below.)

For conventional whole‐cell recordings, used to estimate possible effects of ECM degradation on [Cl^−^]_o_, the pipette‐filling solution contained (in mM): K‐acetate 125, Na‐acetate 3.0, NaCl 6, CaCl_2_ 0.9, Mg‐ATP 5.0, Na_2_‐GTP 0.4, EGTA 2.5, HEPES 10, pH 7.2 (adjusted with KOH).

For cell‐attached recordings used to analyze the effects of presynaptic stimulation on postsynaptic firing frequency, the pipette was filled with standard extracellular solution (see above).

A HCO_3_^−^‐buffered extracellular solution used for some control experiments (Fig. 1*D, E*) contained (in mM): 124 NaCl, 25 NaHCO_3_, 1.1 NaH_2_PO_4_·H2O, 3.5 KCl, 2.0 CaCl_2_, 2.0 MgSO_4_, 10 glucose. The solution was equilibrated with a gas mixture of 95 % O_2_ and 5 % CO_2_.

**Figure 1.**
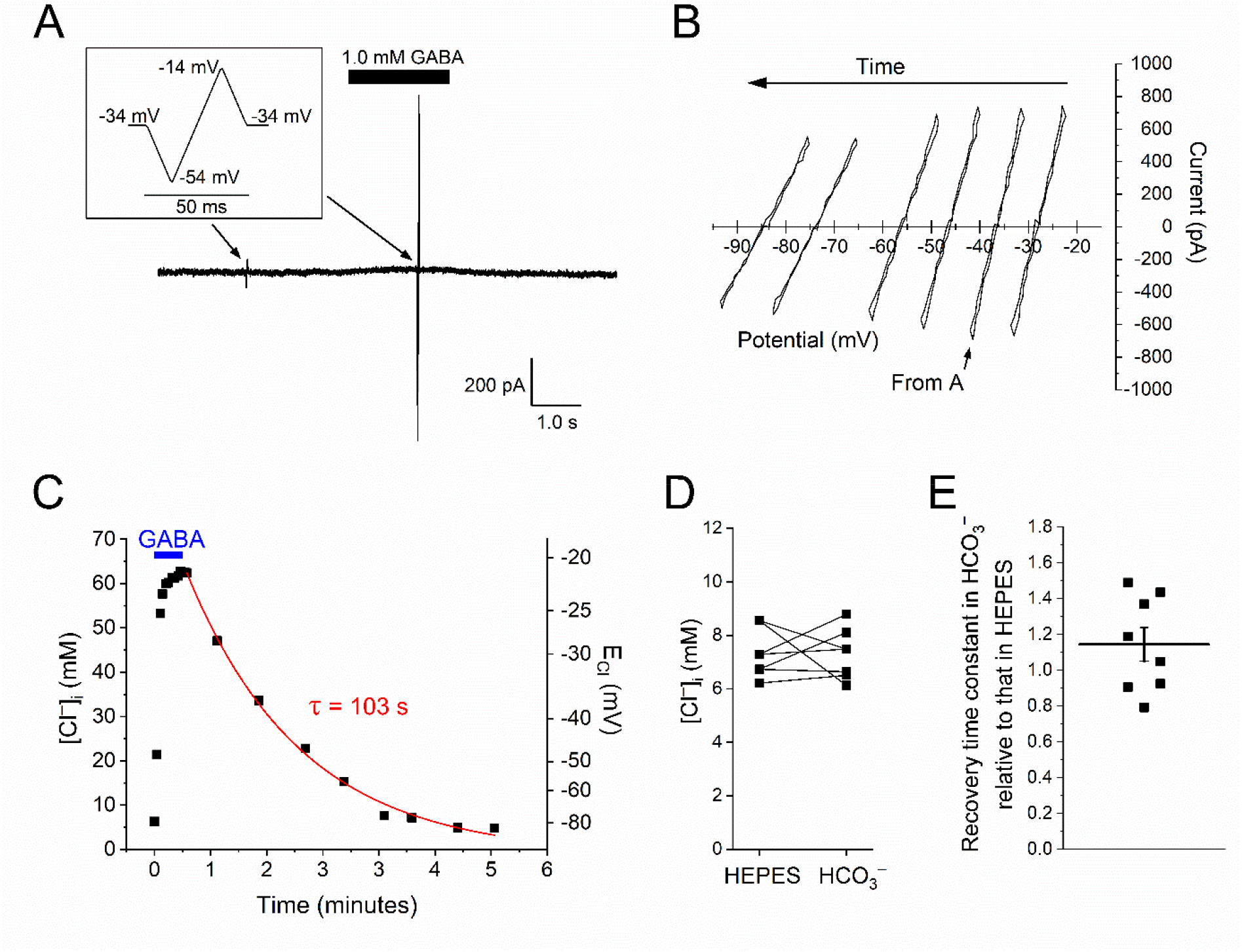
Principle of [Cl^−^]_i_ recovery measurement. ***A***, During and after Cl^−^ loading, voltage ramp sequences (inset) were applied in pairs, without and with GABA, from a holding voltage close to the expected *E*_Cl_. Here, one pair is shown, corresponding to the *I‐V* relation marked “From A” in ***B***. (Cf Fig. 2*A*, for similar current response at a shorter time scale.) ***B***, *I‐V* relations obtained during [Cl^−^]_i_ recovery phase (after correction for series resistance and subtraction of “leak” components; cf Fig. 2*A*). Note the shift in negative direction with time. ***C***, [Cl^−^]_i_ time course during loading (with 1.0 mM GABA, as indicated, at a membrane potential of ~–20 mV) and recovery, computed from reversal potentials (*E*_GABA_). Red curve shows mono‐exponential function fitted to the decay, with time constant indicated. (*E*_Cl_ indicated to the right; note non‐linear scale). ***D***, Basal [Cl^−^]_i_ measured as described above with brain slices in a HEPES‐buffered solution (left data group) and subsequently after 5 ‐ 10 min exposure to a HCO_3_^−^‐buffered solution (same cells; right data group). Cells were kept with current clamped at zero to achieve resting condition, except during measurement. [Cl^−^]_i_ was calculated from the reversal potential of the GABA‐evoked current as described in the main text. The difference between the two solutions was not significant (p = 0.61, paired Wilcoxon signed rank test). ***E***, Ratio between [Cl^−^]_i_ recovery time constants (measured as in ***C***) with a HEPES‐buffered extracellular solution and with a HCO_3_^−^‐buffered solution, in the same cells. The ratio is not significantly different from 1 (p = 0.31, one sample Wilcoxon signed rank test).

The Cl^−^ activity coefficient was calculated using equation 3.14 given by Davies (1962) with z (valency) taken to be 1. (This empiric equation is suitable for ionic strength > 0.1 M and < 0.5 M; see e. g. Butler 1964; Stumm & Morgan, 1996.) For all solutions used for electrophysiology, the Cl^−^ activity coefficient was 0.76; the difference between solutions being ≤ 0.6 %. The activity coefficient for Cl^−^ in the complex intracellular environment during perforated‐patch recording is not known, but may be assumed relatively similar to that in the extracellular solution (Freedman & Hoffman, 1979; Friedman, 2010). The small difference between solutions with different Cl^−^ concentrations is a consequence of Cl^−^activity depending mainly on total ionic strength (Davies, 1962; Friedman, 2010) and justifies the use of concentrations in the Nernst equation.

### Electrophysiological recording

The general techniques and recording equipment used were as previously described (Karlsson et al., 2011). Patch pipettes used had a resistance of 3.5 – 4.2 MΩ when filled with standard pipette‐filling solution and immersed in standard extracellular solution. The gramicidin‐perforated patch method (Abe et al., 1994; Kyrozis and Reichling, 1995) was used, in the majority of experiments, to avoid artificial change of [Cl^−^]_i_ and ionic solutions were chosen to favour currents carried by Cl^−^ (i. e. no HCO_3_ ^−^; see below). Electrophysiological recording in brain slices was performed from neurons located at a depth of 50 – 150 µm, in a majority of cases 75 – 125 µm, and in no case closer to the surface or bottom of the slice than 50 µm. Neurons near the center of the slice were included. Neurons with a resting membrane potential ≤ –45 mV and an input resistance ≥ 100 MΩ were included. All studied cells had a healthy visual appearance according to usual standards (Moyer and Brown, 2002) as judged by microscopic inspection using infrared‐differential interference contrast optics. The slices were continuously perfused by gravity‐fed extracellular solution provided by a custom‐made perfusion pipette positioned 100 – 200 μm from the studied cell, except for experiments with presynaptic stimulation, in which the distance was 1 – 5 mm. Exchange to solution containing GABA, glycine and/or other substances was controlled by solenoid valves via a computer. Liquid‐junction potentials were calculated using the Clampex software (versions 9 & 10; Molecular Devices, CA, USA) and have been subtracted in all potentials given. Treatment of series resistance (*R*_s_) between pipette and cytoplasm is described below.

Unless stated otherwise, recording from cells in slices were made without “cleaning” (see below) and with only a slight positive pressure applied to the patch pipette before sealing, just sufficient to prevent clogging of the pipette tip during penetration of the tissue, with the purpose of minimal effects on the structures beside the membrane patch. In a subset of cells described separately, a cleaning procedure, as described by Edwards et al. (1989) was used with the purpose to systematically remove the ECM covering the exposed surfaces (facing the top and sides of the slice) of the cell. This implied repeated and alternated pressure and suction via a separate, large pipette (tip diameter 5 – 20 µm) filled with standard extracellular solution (see above) (cf Fig 2 in Edwards et al., 1989), before using a regular patch pipette for recording.

**Figure 2.**
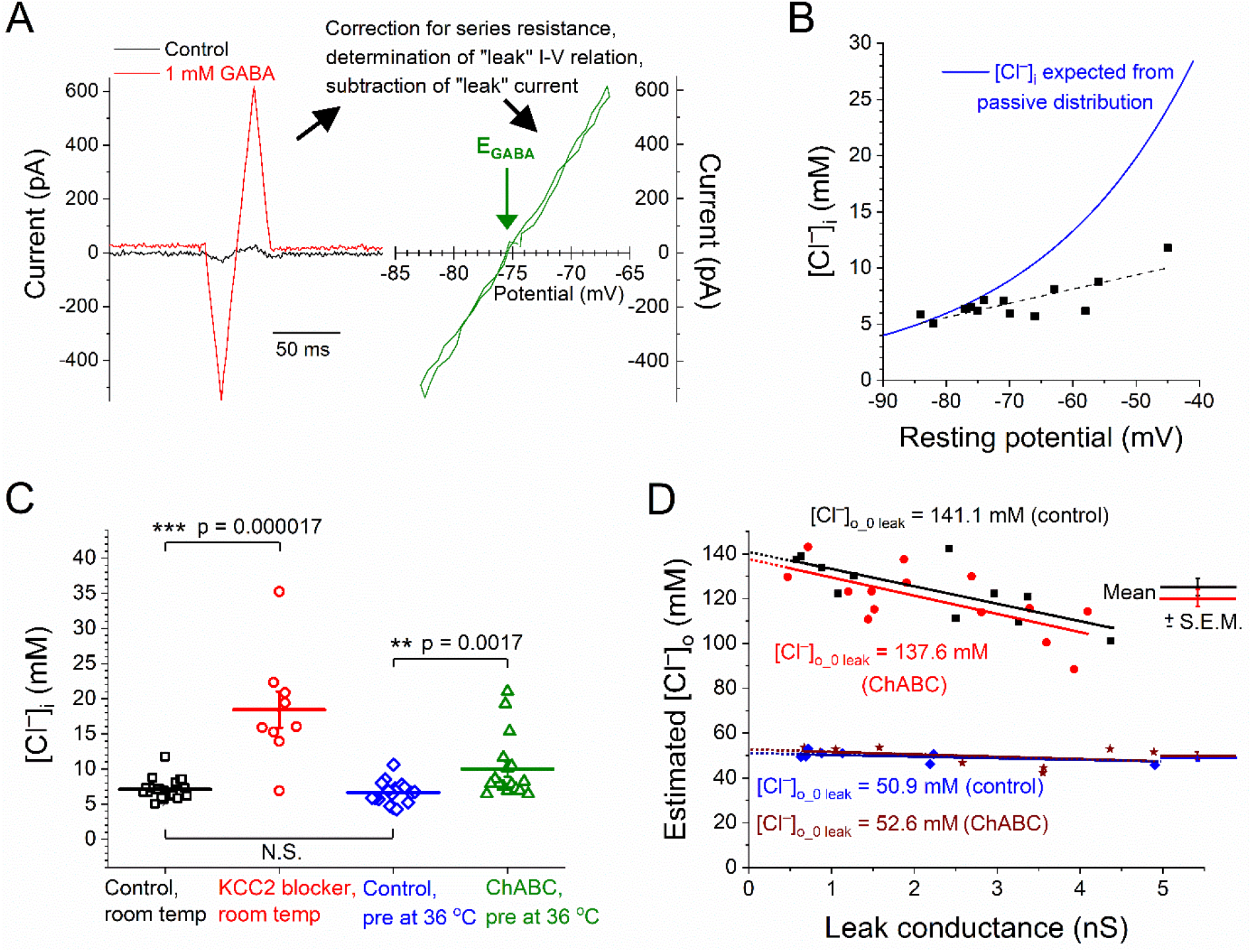
Basal [Cl^−^]_i_ depends on KCC2 and on the ECM, but [Cl^−^]_o_ does not depend on the ECM. ***A***, Raw currents (left) in response to voltage ramps (–74 to –94 to –54 to –74 mV) in the absence or presence of GABA, showing principle of measurement. *I‐V* curve (right) gives *E*_Cl_ (= *E*_GABA_). ***B***, Basal [Cl^−^]_i_ versus resting potential (dashed line: linear fit) in comparison with [Cl^−^]_i_ computed for passive distribution (blue line). ***C***, Significant effects of KCC2 antagonist and of pre‐treatment with ChABC on [Cl^−^]_i_. ***D***, Estimated [Cl^−^]_o_ versus leak conductance, with nominal bulk [Cl^−^]_o_ of 146.4 mM (upper two lines and symbol sets) or 50 mM (lower two lines and symbol sets). Lines: linear fits, dotted extrapolation to zero leak conductance with corresponding [Cl^−^]_o_0 leak_ indicated. Horizontal lines and error bars (in ***C*** and ***D***, right) denote mean ± S.E.M.

#### Measurement of resting potential, calculation of passive [Cl^−^]_i_ distribution and estimation of leak conductance

The resting potential was measured in the current‐clamp mode with zero command current. Similar values were obtained from the reversal potential of currents during test voltage ramps (see below) without agonist in the voltage‐clamp mode, when holding voltage was close to the reversal potential. The [Cl^−^]_i_ expected from a passive distribution was calculated from the Nernst equation using the known [Cl^−^]_o_ and with the equilibrium potential for Cl^−^ (*E*_Cl_) set to the resting potential for each individual cell. The leak conductance was estimated from the slope of a linear fit to the portion of *I‐V* relation around (± 5 mV) the zero current voltage during voltage ramps (± 1.6 V s^‐1^) in the absence of ligands.

#### Measurement of E_GABA/glycine_ and E_Cl_ by use of voltage ramps

The reversal potential for currents evoked by GABA or glycine (*E*_GABA_ and *E*_glycine_, respectively) was assumed to be described by the equation *E*_GABA/glycine_ = R T F^−1^ ln (([Cl^−^]_i_ + P_rel_ [HCO_3_^−^]_i_) ([Cl^−^]_o_ + P_rel_ [HCO_3_^−^]_o_)^−1^) where P_rel_ is HCO_3_^−^/Cl^−^ permeability ratio of GABA or glycine receptors and R, T and F denote the gas constant, absolute temperature and Faraday constant, respectively. P_rel_ was assumed to be 0.2 (Bormann et al., 1987). The gramicidin pores are highly permeable to H^+^ (Myers and Haydon, 1972) and thus intracellular pH may be assumed equal to the pH of the pipette‐filling solution. The intracellular [HCO_3_^−^] calculated by the Henderson‐Hasselbalch equation with CO_2_ partial pressure from Boron (2009a), solubility constant and p*K* value, from Boron (2009b) or Kajino et al. (1982), is ~ 0.1 mM. This implies a negligible effect of HCO_3_ ^−^ on the reversal potential for currents through GABA_A_ receptors (*E*_GABA_) with the standard HEPES‐buffered extracellular solution (see above) used in most experiments, where *E*_Cl_= *E*_GABA_ = *E*_glycine_. (In the absence of HCO_3_ ^−^ in the recording pipette, *E*_GABA_ fits well with the Cl^−^ equilibrium potential, *E*_Cl_; Chavas and Marty, 2003).

The reversal potential was obtained from *I‐V* relations constructed from currents recorded during voltage ramps, given at a rate of ± 1.6 V s^−1^ (Karlsson et al., 2011). A ramp sequence consisting of negative and positive voltage ramps as shown in Fig. 1A (inset) was applied twice, in the absence and presence of GABA or glycine, for recording of “leak” and GABA/glycine‐evoked (plus leak) currents, respectively. (The timing of the second ramp was usually in the mid‐50% portion of the 2‐s long GABA/glycine application.) Occasionally unclamped action potentials were generated during the rising phase of voltage ramps in control solution without GABA/glycine. Portions of current with action potentials were excluded from further analysis. The holding voltage (V_h_) before, after and in‐ between ramp sequences depended on the type of measurement: For estimation of basal [Cl^−^]_i_, V_h_ was kept close to the resting potential. To follow recovery of [Cl^−^]_i_ after loading to high levels (see below), V_h_ was kept close to the expected *E*_Cl_ (as estimated in preliminary experiments, see below), to minimize Cl^−^ current (cf the small GABA‐evoked current at holding potential, Fig. 1*A*). Influence on [Cl^−^]_i_ was further minimized by using a symmetrical voltage ramp protocol (Yelhekar et al., 2016) (Fig. 1*A*, inset), yielding approximately similar Cl^−^ flux in inward and outward directions (Fig. 2*A*). For measurement of basal [Cl^−^]_i_, special care was taken to avoid changes in [Cl^−^]_i_ during seal formation between patch pipette and cell membrane: The current‐clamp recording mode with *I* = 0 was used during the gramicidin perforation, except for brief tests of *R*_s_. (This implies that no current from the recording pipette may change the membrane potential during the seal formation and the parallel gramicidin perforation of the membrane.) In addition to these precautions, we quantified the effect of the [Cl^−^]_i_‐probing GABA/glycine application and voltage ramps on [Cl^−^]_i_ as described below.

#### Procedure to minimize conductive (passive) Cl^−^ flux during recovery from high [Cl^−^]_i_

To minimize conductive (passive) Cl^−^ flux for better estimating the time course of transporter‐mediated changes in [Cl^−^]_i_, the holding voltage was kept as close as possible to the variable *E*_Cl_ during recovery from high [Cl^−^]_i_. For this, an approximate time course of *E*_Cl_ was first obtained from test runs, with initial holding voltage at an estimated (assumed) *E*_Cl_: The magnitude and polarity of the first current response to GABA/glycine, delivered shortly after the start of recovery period (end of GABA/glycine application for loading) gave information on the magnitude and polarity of the difference between estimated and true *E*_Cl_. If the difference was large, the test run was aborted. If the difference was small, the test run was continued with a second pair (for leak and GABA/glycine‐evoked currents) of ramp sequences from a new holding voltage. Again, the difference between estimated and true *E*_Cl_ was evaluated for decision on continuation or abortion of the test run. In the absence of large deviations between estimated and true *E*_Cl_, the test run was continued with further ramp sequences until recovery was near complete / a stable *E*_Cl_ was reached.

After a few test runs for each cell and condition, it was possible to predict the approximate time course of *E*_Cl_ recovery, so that holding voltages reasonably close to *E*_Cl_ could be used. Note, however, that unless the holding voltage (estimated *E*_Cl_) exactly matched the true *E*_Cl_, the GABA/glycine application will result in some positive or negative current. It was important, therefore, to estimate the influence of the GABA/glycine applications on [Cl^−^]_i_, as described below. The lack of correlation between [Cl^−^]_i_ recovery time course and leak conductance (see Results) confirmed that influence of passive Cl^−^ flux was of no major importance for the recovery.

#### Influence of measurement procedure on [Cl^−^]_i_

The influence of test pulses (with GABA or glycine application) on [Cl^−^]_i_ was estimated on basis on recorded charge transfer and the estimated cytosolic volume in which Cl^−^ equilibrates. This “equivalent volume” was obtained from the GABA‐evoked charge transfer during Cl^−^ loading (see below) and the measured rapid large change in [Cl^−^]_i_ between two test ramps during the loading phase. In dissociated neurons, for which cell volume is easily estimated, this “equivalent volume” corresponds to about half the cell volume (Karlsson et al., 2011), in good agreement with expected cytosolic volume. The charge transfer associated with a test pulse of GABA during [Cl^−^]_i_ recovery was on average expected to change [Cl^−^]_i_ by 0.90 ± 0.48 mM (mean ± SEM; 8 cells).

#### Loading cells with Cl^−^

Loading cells to a high [Cl^−^]_i_, for enabling subsequent estimates of Cl^−^ transport capacity, was made by combining a depolarized holding voltage (~ –20 ‐ 0 mV) with application of 1.0 mM GABA (Sigma‐Aldrich) or 1.0 mM glycine (Sigma‐Aldrich) for 20 ‐ 30 s (Karlsson et al., 2011). A single voltage‐ramp sequence, applied before the agonist, was then used to estimate leak currents for repeated [Cl^−^]_i_‐probing voltage ramps during this long‐lasting agonist application (cf Fig. 2A‐C in Yelhekar et al. 2017).

#### Subtraction of leak and capacitive currents

The *I‐V* relation for “leak” currents induced by the voltage ramps in the absence of ligands for Cl^−^ permeable channels was (after correction for R_s_ effects as described below) often well fitted by a linear function. In some cases, especially at relatively positive voltages, a slightly curved relation was obtained, likely due to some contribution of voltage‐sensitive currents. In the latter cases, the *I‐V* relation was well fitted by the exponential function: *I* = A + B *e*^*V*/C^ (A, B and C being constants). The obtained “leak” *I‐V* relation was used to calculate and subtract leak (and small voltage‐gated) current components from the current measured in the presence of GABA or glycine during short ligand application (Karlsson et al., 2011; Yelhekar et al., 2017). The use of the mathematically expressed relation between leak current and *R*_s_‐ corrected voltage rather than the current directly recorded during the voltage ramps (in the absence of GABA/glycine) provides a better estimate of leak current during voltage ramps in the presence of GABA/glycine, since the voltage with and without agonist as a rule differs as a consequence of the induced currents across the series resistance (*R*_s_).

The current during the rising as well as declining phases of the voltage ramp sequence (Fig. 1*A*, inset) was used for the *I‐V* relation (apparently seen as two superimposed curves, e.g. Fig. 1*B*). Often, the reversal potential from the rising and declining phases of the ramp sequence were not significantly different (Fig. 1*B*; see also Fig. 2*A*, right), but in some cases a contribution of capacitive current (which is constant but of opposite sign during rising and declining phases) caused a slight offset of the two curves. In the latter cases, *E*_Cl_ (or *E*_GABA_ or *E*_glycine_) was obtained from the midpoint (mean) between the two reversal potentials. [Cl^−^]_i_ or [Cl^−^]_o_, depending on the experiment, was subsequently calculated from the relation between reversal potential and [Cl^−^] given above (for *E*_Cl_, Nernst equation). As previously described (Karlsson et al., 2011), GABA_B_ receptors do not contribute significantly to GABA‐evoked currents in the studied neurons.

#### Series resistance compensation

Online series resistance (*R*_s_) compensation during electrical recording was not used since it is usually accompanied by additional noise and since full compensation is not possible with maintained electrical stability (see e.g. The Axon Guide. Electrophysiology & Biophysics Laboratory Techniques”, 3^rd^ ed., 2012, Molecular Devices, CA, U.S.A.). To better account for effects of *R*_s_, all voltages were corrected offline by subtracting *R*_s_ *I*, where *I* is raw recorded current and *R*_s_ the series resistance as measured by the “membrane test” procedure in the Clampex software. The above description of leak and capacitive current subtraction explains how the different effects of *R*_s_ on membrane voltage during recording of leak currents compared with agonist‐induced currents were taken into account. Series resistance in the gramicidin‐perforated patch mode was 37 ± 2 MΩ (n = 110) and in the whole‐cell mode 23 ± 2 MΩ (n = 25). For recording of voltage‐gated Ca^2+^ currents in amphotericin‐perforated patch mode, see below.

#### Whole‐cell recording for estimation of E_Cl_

The whole‐cell recording mode was used in some experiments to dialyze the cytoplasm with a known Cl^−^ concentration and enable estimation of the [Cl^−^]_o_ that contributes to *E*_Cl_, and reveal a hypothetical difference from [Cl^−^] in the external bulk solution as suggested by Glykys et al. (2014a). In these experiments we used Cs^+^ rather than K^+^ as the major cation in the pipette, to reduce background noise, to improve voltage‐clamp control (Chen et al., 1999) and to reduce Cl^−^ transport by KCC2 (Cs^+^ competing with other cations; inhibition constant 9.7 mM; Williams and Payne, 2004). In the whole‐cell mode, equilibration of Cl^−^ between pipette and cytosol is expected within a few seconds for spherical cells (Pusch and Neher, 1988), but deviations of [Cl^−^]_i_ from [Cl^−^]_pip_ may possibly arise as a consequence of influence of dendrites or Cl^−^ leak across the membrane (or across pipette‐membrane seal). Such influence was here suggested by the correlation between leak conductance and measured *E*_Cl_ (and thus calculated [Cl^−^]_o_) when high [Cl^−^] was present in the external bulk solution (Fig. 2*D*, upper two lines). With reduced [Cl^−^] in the external bulk solution, however, the observed data (Fig. 2*D*, lower two lines) did not reveal any significant influence of such leak.

#### Amphotericin B‐perforated recording of Ca^2+^ currents

Whole‐cell Ca^2+^ currents were recorded in the amphotericin B‐perforated patch configuration (120 µg ml^−1^ amphotericin B; Sigma‐Aldrich) under voltage‐clamp conditions, with voltage‐gated Na^+^ currents blocked by 2.0 µM tetrodotoxin (Latoxan) in the extracellular solution and voltage‐gated K^+^ currents blocked or reduced by Cs^+^ substituted for K^+^ in the pipette solution. Currents were activated by voltage steps from –54 mV to – 14 mV. The leak currents and capacitive currents were subtracted using scaled current responses to negative potential steps. The finding of linear *I‐V* relations for currents evoked in preoptic neurons in response to voltage steps in the range –40 to –140 mV justified this procedure (Sundgren‐Andersson and Johansson, 1998). Series resistance compensation was not used for these recordings and maximum voltage error (at peak current) was estimated to 15 mV. (Series resistance in the amphotericin‐perforated patch mode was 25 ± 2 MΩ; n = 19.) Further, only the relative components of Ca^2+^ current amplitudes and charge transfer in each cell were used for analysis, to account for intercellular variability in absolute Ca^2+^ currents.

#### Presynaptic stimulation

Extracellular stimulation of presynaptic nerve fibers was made as previously described (Malinina et al., 2005). Postsynaptic responses to such stimulation were recorded in the cell‐attached mode (10 – 18 cells, depending on type of experiment) and verified in the gramicidin‐perforated patch mode (see above; 3 – 4 cells for each type of experiment). Effects on firing frequency were analyzed with data pooled from the two used recording configurations.

All electrical recordings were performed at room temperature (21 – 23 °C).

### Protein estimation

Brain slices from the medial preoptic area (as described above) were used for protein estimation by Western blot (see below). From each animal, a maximum of three adjacent brain slices were used, and divided along the midline to achieve a maximum of six hemispheric slices for separate exposure to different test substances or use as control. The choice of (left or right) hemispheric slice for the different exposures was made at random to avoid any systematic inter‐hemispheric differences. The protein samples obtained from the brain tissue was split and run repeatedly (three times each, with average taken, for improved reliability) on separate gels.

#### Estimation of total KCC2 protein and of αII‐spectrin fragments

For total protein extraction, the brain slices were homogenized in RIPA buffer (Thermo Fisher) containing a protease inhibitor cocktail (Sigma‐Aldrich). In experiments for αII‐spectrin degradation, 1.0 % of each phosphatase inhibitor cocktail 2 and 3 (Sigma‐Aldrich) as well as EDTA and EGTA (5.0 mM each; Sigma‐Aldrich) were added to prevent additional lysis‐induced Ca^2+^ dependent calpain activation. Non‐soluble proteins were separated by centrifugation at 13 000 g for 10 min at 4 °C. The protein concentration of individual samples was measured using the BCA protein assay kit (Thermo Fisher). Protein samples (8 – 12 μg; 30 µg for αII‐spectrin degradation assessment) in reducing (5.0 % 2‐mercaptoethanol) sample buffer (Bio‐Rad) were heated at 70° for 30 or 10 (for αII‐spectrin) min and loaded on precast 4 – 12 % polyacrylamide gels (Bio‐Rad) or on homemade 8 % gels (for αII‐spectrin) and separated by SDS‐PAGE electrophoresis. Subsequently, proteins were transferred to polyvinylidene difluoride (PVDF) membranes (Thermo Fisher) in freshly prepared buffer containing 25 mM Tris‐base, 192 mM glycine and 20 % (v/v) methanol. Non‐specific antibody binding was blocked by 5.0 % blotting‐grade blocker non‐fat dry milk (Bio‐Rad) in Tris‐buffered saline (TBS) and tween 20 (0.1 %) for 1 h at room temperature. Membranes were incubated with primary antibodies in 5.0 % non‐fat dry milk‐ containing TBS‐tween solution. Primary antibodies, anti‐KCC2 (1: 5000; Millipore), anti‐transferrin receptor (1:500; Invitrogen), anti‐α‐fodrin (1:5000; Enzo, clone AA6) and anti‐β‐tubulin (1:10000; Abcam) were used to stain KCC2, transferrin receptor, αII‐spectrin (full form and fragments) and β‐ tubulin respectively. After overnight incubation with primary antibodies, membranes were incubated with horseradish peroxidase (HRP)‐linked anti‐mouse (1:5000; GE Healthcare) or HRP‐linked anti‐ rabbit (1:1000; Cell signaling) secondary antibodies at room temperature for 1 h. Chemiluminescent detection was done using Luminata Forte Western HRP substrate (Millipore) and images were taken on an Odyssey Fc imaging system (LI‐COR). Quantification of chemiluminescent signals was made using Image studio (version 3.1, LI‐COR). Protein levels of KCC2, transferrin receptor and αII‐spectrin (full form and fragments) were estimated relative to the level of β‐tubulin.

#### Estimation of KCC2 dimer and monomer levels in membrane fraction

ChABC‐ or vehicle‐ treated brain slices (as described above) were homogenized in lysis buffer containing 50 mM Tris·HCl, pH 7.4, 1.0 mM EDTA, supplemented with 1.0 % CLAP (chymostatin (Sigma‐Aldrich), leupeptin hemisulfate, antipain‐dihydrochloride, pepstatin A (Carl Roth)), 0.5 mM phenylmethyl sulphonyl fluoride (PMSF; Carl Roth), 1.0 % each phosphatase inhibitor cocktails 2 and 3 (Sigma‐Aldrich), and 25 mM iodoacetamide (Sigma‐Aldrich), used to block free thiol groups and prevent ex‐vivo KCC2 multimerization (Blaesse et al., 2006; Mahadevan et al., 2014). To isolate membrane proteins from cytosolic ones, homogenized samples were centrifuged at 10 000 g for 30 min. The obtained pellet was resuspended in CHAPS buffer (50 mM Tris·HCl, pH 7.4, 10 mM CHAPS (Glycon Biochemicals), 0.05 mM EDTA) containing CLAP, PMSF, phosphatase inhibitor cocktails 2 and 3 and iodoacetamide in concentrations as above and allowed to solubilize for 3 h on a rotating platform at 4 °C. The samples were centrifuged for 1 h at 10 000 g and the supernatants were recovered. Cleared lysates (10 µg) were suspended in non‐reducing electrophoresis sample buffer (6.0 % SDS, 20 % glycerol, 62.5 mM Tris·HCl, pH 6.8, 0.01 % bromophenol blue) to prevent disassembly of KCC2 oligomers, which are sensitive to sulfhydryl‐reducing agents but resistant to SDS (Blaesse et al., 2006). Subsequently, the proteins were separated by SDS‐PAGE on 7.5 % gels and immunoblotted with anti‐ KCC2 (1:1000) and anti‐calnexin antibody (1:2000; Enzo Life Sciences). Densitometric analysis of corresponding bands was performed by Image Studio software (LI‐COR Biosciences). KCC2 protein level in the membrane fraction was estimated relative to the level of the membrane protein calnexin.

### Experimental design and statistical analysis

Statistical analysis was performed using OriginPro, versions 9.0 – 2019. All data are presented as mean ± SEM unless otherwise stated. Effects were evaluated by the non‐parametric one‐sample Wilcoxon test, paired Wilcoxon signed rank test or Mann‐Whitney test, as indicated. Spearman′s rank correlation coefficient was used to evaluate possible correlations between measured variables or recording parameters. For electrophysiological experiments, including estimates of Cl^−^ concentrations, numbers (n) given denote the number of cells studied, with paired tests used for comparison of different conditions for the same cell. The cells used to analyse each test (substance/condition) were obtained from a minimum of five (and up to 22) brain slices from a minimum of four (and up to 19) rats. For Western blot data, numbers (n) denote the number of animals studied, with paired tests used to compare differently treated brain tissue from right and left hemispheres of the same individual. P‐values < 0.05 were taken to indicate significant differences.

## Results

### The low basal [Cl^−^]_i_ depends on KCC2 as well as on the ECM

We estimated basal [Cl^−^]_i_ in preoptic neurons in slice preparations of rat brain, by direct recording of Cl^−^ currents through GABA_A_‐ or glycine‐receptors during voltage ramps (Karlsson et al., 2011) (Fig. 2*A*). The gramicidin‐perforated patch method (Abe et al., 1994; Kyrozis and Reichling, 1995) was used to avoid artificial change of [Cl^−^]_i_ and ionic solutions were chosen to favour currents carried by Cl^−^ (i. e. no HCO_3_ ^−^; see Materials and Methods). In control conditions, [Cl^−^]_i_ was significantly lower than expected from a passive distribution (7.0 ± 0.5 mM vs 10.2 ± 0.5 mM; n = 13, p = 0.0024, paired Wilcoxon signed rank test; Fig. 2*B*). Such a difference is expected as a consequence of KCC2‐mediated transport, with the effect on [Cl^−^]_i_ depending on the membrane Cl^−^ conductance (Johansson et al., 2016). Here, the selective KCC2‐antagonist VU0255011‐1 (10 µM for 30 min) increased [Cl^−^]_i_ to 18.4 ± 2.6 mM (n = 9, p = 1.7 10^−5^, Mann‐Whitney test; Fig. 2*C*) without affecting the resting potential (control: –69 ± 3 mV, n = 13; VU0255011‐1: –68 ± 3 mV, n = 9; p = 0.59, Mann‐Whitney test between groups), showing that KCC2 is critically required for the low basal [Cl^−^]_i_.

We next examined if the ECM is important for the basal [Cl^−^]_i_. Chondroitinase ABC treatment (0.2 U ml^−1^; see Materials and Methods) resulted in ECM degradation, as verified histochemically by WFA staining (Fig. 3). In parallel, basal [Cl^−^]_i_ increased from 6.6 ± 0.4 mM (n = 18) to 10.0 ± 1.1 mM (n = 17, p = 1.7 10^−3^, Mann‐Whitney test; Fig. 2*C*). Basal [Cl^−^]_i_ did not significantly depend on whether a HEPES‐ or bicarbonate‐buffered extracellular solution was used (Fig. 1*D*). The resting potential did not differ significantly (Mann‐Whitney test, p = 0.30) between control cells (–70 ± 2 mV, n = 19; exposed to the same temperature as ChABC‐treated cells) and those exposed to ChABC (–67 ± 2 mV, n = 20).

**Figure 3.**
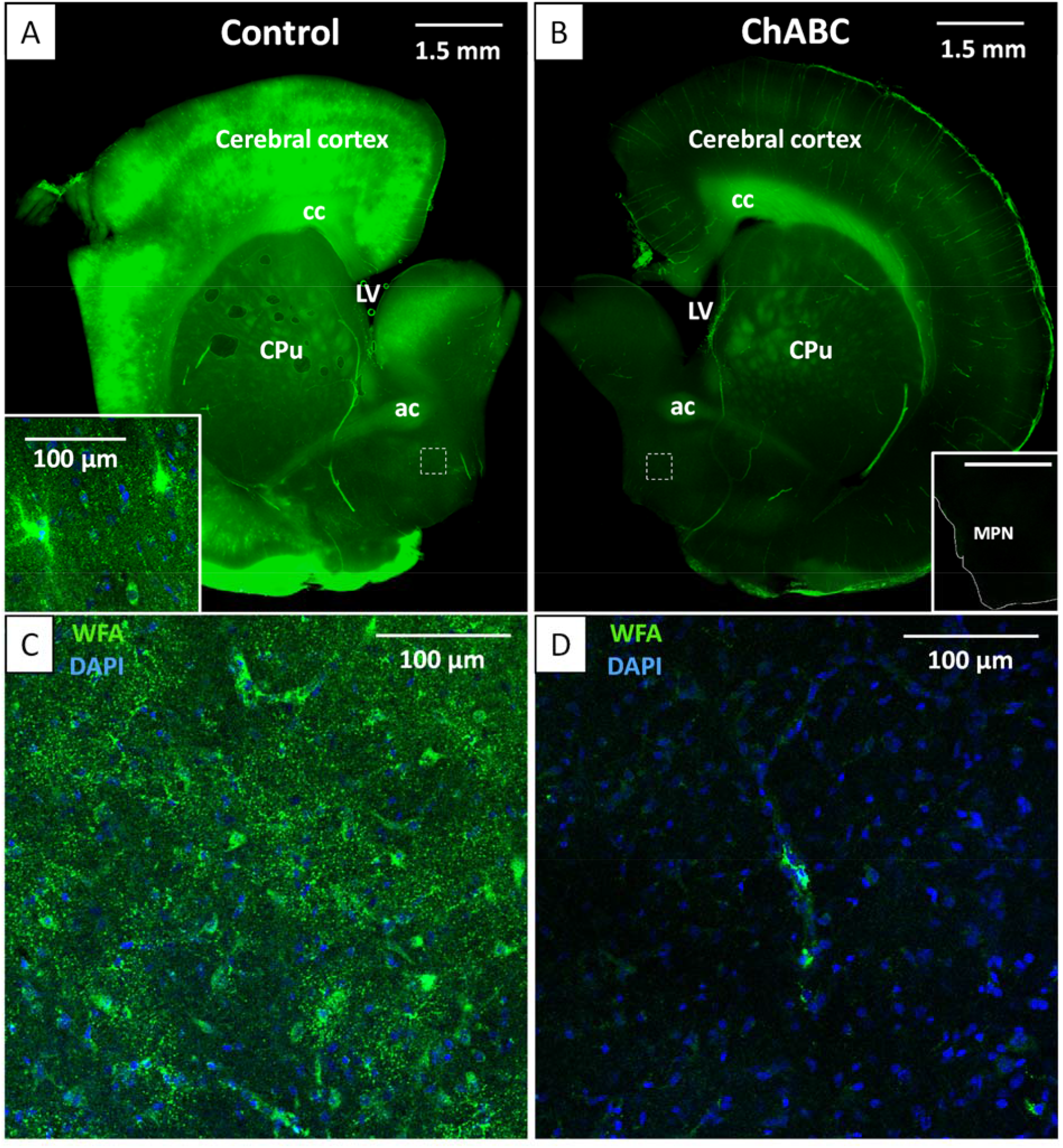
ChABC treatment degrades ECM. ***A, B***, Low‐magnification images of hemispheral coronal brain slices incubated for 2 h in vehicle (0.1 % BSA, Control, ***A***) or ChABC (0.2 U ml^−1^, ***B***) and stained with *Wisteria floribunda* agglutinin (WFA) to label chondroitin sulphate proteoglycans present in the ECM. Dashed white boxes denote the approximate location of the medial preoptic nucleus (MPN) shown at higher magnification below (***C, D***). Note, that the WFA signal is much weaker in the slice treated with ChABC as compared with the control. Inset in ***A*** shows a confocal image from neocortex (control), obtained with 20× objective, with typical WFA‐positive perineuronal nets surrounding some cells. Inset in ***B*** shows an area including the MPN from a negative control slice, incubated without WFA (scale bar 1.5 mm as in ***A***; thin white line drawn to show border of slice). ***C, D***, Confocal images obtained with 20× objective; control (***C***) and ChABC treated (***D***) medial preoptic area; merged WFA (green) and DAPI (blue; for DNA in cell nuclei) staining. cc – corpus callsosum, ac – anterior commissure, CPu – caudate putamen (striatum), LV – lateral ventricle.

### ECM degradation does not affect local [Cl^−^]_o_

According to Glykys et al. (2014a), ChABC increases [Cl^−^]_i_ as a consequence of removed ECM charges and the associated increase in local [Cl^−^]_o_. However, the idea that local extracellular charges would alter the driving force for Cl^−^ across the cell membrane has been questioned on basis of thermodynamic principles (Kaila et al., 2014; Voipio et al., 2014) and such effects may possibly occur only if there is a local displacement of Cl^−^ in the immediate vicinity of the Cl^−^‐permeable channels (Savtchenko et al., 2017). To experimentally verify that the driving force for Cl^−^ is not affected by ECM charges, we estimated whether the [Cl^−^]_o_ contributing to the driving force for Cl^−^, possibly different from the Cl^−^ concentration in the external bulk solution ([Cl^−^]_o_bulk_), was affected by ECM degradation. Therefore, we used the whole‐cell patch‐clamp mode to dialyze the cell with a known Cl^−^ concentration (7.8 mM), which enabled estimation of [Cl^−^]_o_ from *E*_GABA_, obtained with voltage ramps as above, and from the defined [Cl^−^]_i_. In these experiments, Cs^+^ was substituting K^+^ as major cation in the pipette, to reduce KCC2‐mediated transport (Williams and Payne, 2004; see Materials and Methods). With [Cl^−^]_o_bulk_ = 146.4 mM, the estimated [Cl^−^]_o_ was 125 ± 4 mM (n = 11), substantially less than [Cl^−^]_o_bulk_. However, the method may be influenced by Cl^−^ leak into the cell, as suggested by a negative correlation between membrane leak conductance and estimated [Cl^−^]_o_ (Fig. 2*D*, black). To improve the estimation, we extrapolated [Cl^−^]_o_ linearly to zero leak, thus obtaining [Cl^−^]_o_ = 141.1 mM, i. e. relatively close to [Cl^−^]_o_bulk_. We next tested the hypothesis that ChABC treatment is accompanied by increased [Cl^−^]_o_. Mean [Cl^−^]_o_ after ChABC treatment was 120 ± 4 mM (n = 14) (Fig. 2*D*, red), not significantly different from control (Mann‐Whitney test), and the improved [Cl^−^]_o_ estimate obtained by extrapolation to zero leak was 137.6 mM. We also estimated [Cl^−^]_o_ when [Cl^−^]_o_bulk_ was 50 mM and less Cl^−^ leak into the cell was expected. In this case, mean [Cl^−^]_o_ was within 1.0 mM from the expected value, in control conditions (49.1 ± 0.9 mM; n = 9; Fig. 2*D*, blue) as well as after ChABC treatment (49.9 ± 1.4 mM; n = 9; Fig. 2*D*, wine). Again, there was no significant difference between control and ChABC‐treated cells (Mann‐Whitney test). The lack of change in measured *E*_GABA_ and estimated [Cl^−^]_o_ contradicts the idea that ChABC increases the driving force by increasing local [Cl^−^]_o_ and thus also contradicts the idea that such effects would be the cause for increased [Cl^−^]_i_ after ChABC treatment.

### KCC2‐mediated [Cl^−^]_i_ recovery rate depends on ECM integrity

From the above results, it seems likely that KCC2 function depends on ECM integrity. To test this hypothesis, we first estimated Cl^−^ transport rate from repeated measurements of [Cl^−^]_i_ during recovery after loading to [Cl^−^]_i_ ~40 ‐ 75 mM (see Materials and Methods and Fig. 1; cf Yelhekar et al., 2017). The recovery showed an exponential time course (time constant 111 ± 14 s; n = 19) and was not significantly altered by the NKCC1‐antagonist bumetanide (10 µM; Sigma‐Aldrich; Fig. 4*A;* n = 8, paired Wilcoxon signed rank test), but was dramatically slowed down by VU0255011‐1 (10 µM; Fig. 4*A, B*). After 14 minutes of recovery, estimated [Cl^−^]_i_ was 5.3 ± 0.3 mM in control solution, but 41.5 ± 2.3 mM in the presence of VU0255011‐1 (p = 7.8 10^−3^, paired Wilcoxon signed rank test; Fig. 4*B*), suggesting a main role of KCC2 for [Cl^−^]_i_ recovery. In similarity with basal [Cl^−^]_i_, the [Cl^−^]_i_ recovery time constant did not significantly depend on whether a HEPES‐ or bicarbonate‐buffered extracellular solution was used (Fig. 1*E*). We subsequently tested the hypothesis that the ECM is important not only for basal [Cl^−^]_i_, but also for the rate of KCC2‐mediated Cl^−^ transport. The [Cl^−^]_i_ recovery time constant, after Cl^−^‐ loading cells in control slices pre‐incubated at ~ 36 °C for 2 h (to enable comparison with ChABC‐treated slices), was 137 ± 18 s (n = 19), not statistically different (Mann‐Whitney test) from that in control slices kept at room temperature (21 – 23 °C). (Note that even when pre‐incubation at 36 °C was used, evaluation of [Cl^−^]_i_ recovery time constant was made at room temperature.) However, ChABC treatment dramatically increased the [Cl^−^]_i_ recovery time constant to 441 ± 84 s (n = 15, p = 4.8 10^−6^, Mann‐Whitney test; Fig. 4*C, D*), confirming that the ECM is important for the rate of KCC2‐mediated Cl^−^ transport.

**Figure 4.**
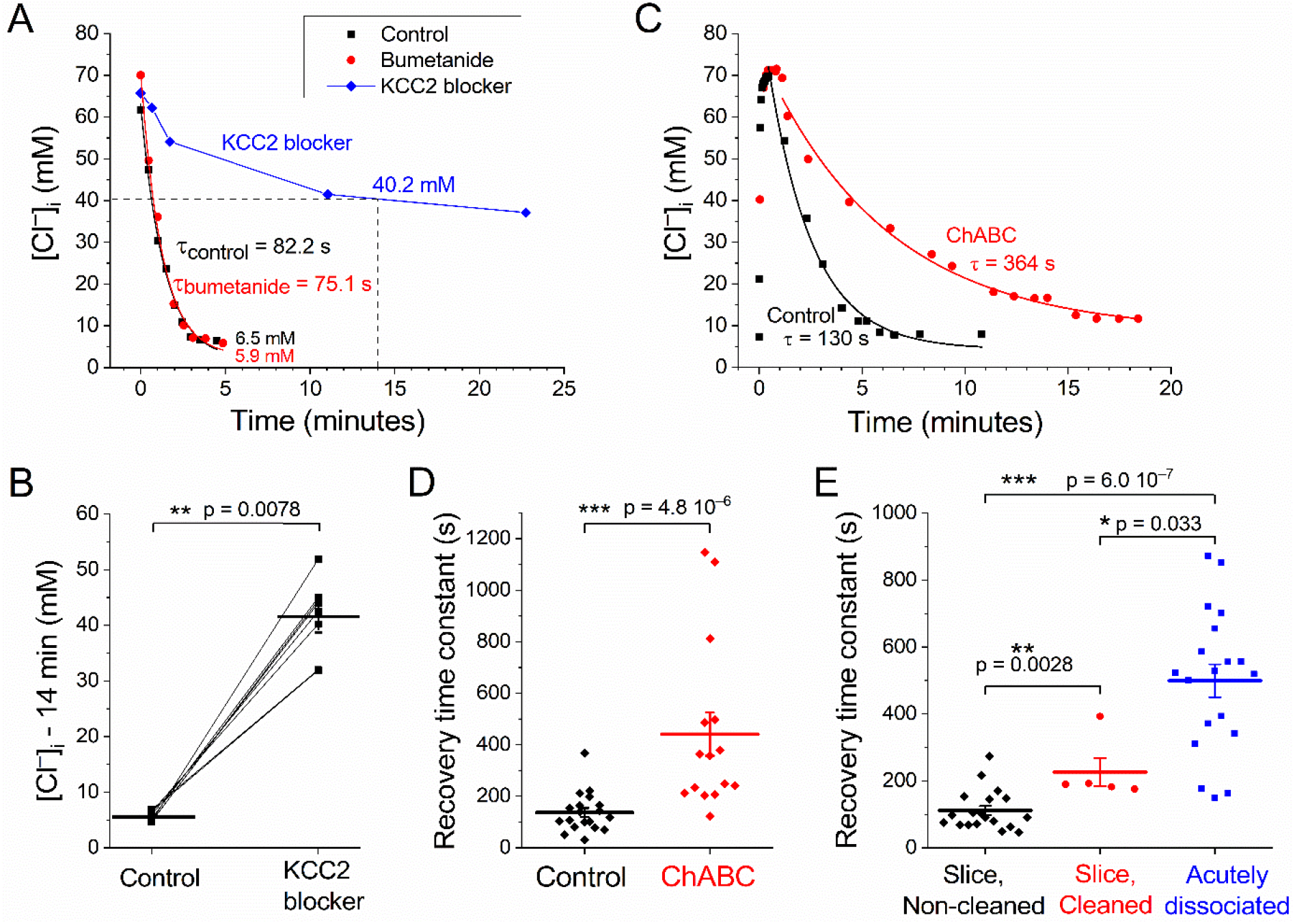
Cl^−^ extrusion rate depends on KCC2 and on the ECM. ***A***, Time course of [Cl^−^]_i_ recovery after load to ~70 mM. Single neuron in control solution, with bumetanide (10 μM) and with VU0255011‐1 (10 μM). Solid lines: exponential fits (control and bumetanide), linear interpolation (KCC2 antagonist). ***B***, [Cl^−^]_i_ after nominally 14 minutes recovery (see below; cf dashed lines in ***A***); control and with 10 μM VU0255011‐1 (paired data from 8 neurons). ***C***, [Cl^−^]_i_ recovery time course after pre‐incubation for 2 h at 35‐37 °C in the absence (Control) or presence of ChABC. ***D***, Significant effect of ChABC on [Cl^−^]_i_ recovery time constants (cf ***C***). ***E***, Significant effect of cell cleaning and of dissociation on [Cl^−^]_i_ recovery time constants (cf ***A*** and ***C***). Data for acutely dissociated cells from Yelhekar et al. (2017). Horizontal lines and error bars (in ***B, D*** and ***E***) denote mean ± S.E.M. The [Cl^−^]_i_ at a nominal time point of 14 min in ***B*** was estimated by extrapolation of the near‐steady [Cl^−^]_i_ in cases where [Cl^−^]_i_ decayed to < 10 mM within 10 minutes and by linear interpolation of [Cl^−^]_i_ at an interval where [Cl^−^]_i_ changed less than ~1 mM/min in cases with slower decay, as illustrated by dashed lines in ***A***.

As an alternative to ChABC treatment, we used the “cleaning” method for patch‐clamp recording in slices (Edwards et al., 1989; see Materials and Methods), with a stream of saline to remove the ECM covering the top and other exposed surfaces of neuronal soma and basal neurites. This type of cleaning is, like attenuation of ECM with β‐D‐xyloside, a procedure that facilitates giga‐seal formation at patch‐clamp recording (Izu and Sachs, 1991). [Cl^−^]_i_ recovery in cleaned cells was much slower than in control conditions (time constant 227 ± 42 s; n = 5, cleaned, vs 111 ± 14, control, n = 19, p = 0.0028, Mann‐Whitney test; Fig. 4*E*). Thus, the two independent tests, with different procedures to affect the ECM, suggest a role of the ECM in controlling the rate of Cl^−^ extrusion. [Cl^−^]_i_ recovery measured previously with similar methods in acutely dissociated preoptic neuronal cell bodies, where we expect even less ECM, is still slower (499 ± 49 s, n = 19; Yelhekar et al., 2017; comparison with present control values: p = 6.0 10^‐7^, Mann‐Whitney test), further strengthening this idea (Fig. 4*E*).

The resting membrane properties of the cleaned cells in slices and those of the acutely dissociated cells differed somewhat from those reported for non‐cleaned cells in slices reported above. Thus, the resting potential was –62 ± 6 mV (n = 5) and –60 ± 3 mV (n = 19) for cleaned and for dissociated cells, respectively. The corresponding values for leak conductance was 2.0 ± 0.3 nS (cleaned) and 0.91 ± 0.15 nS (dissociated) and for membrane capacitance 11 ± 1 pF (cleaned) and 5.5 ± 0.3 pF (dissociated) in the same cell groups. (Series resistance did not differ significantly between any of the studied cell groups). A correlation analysis performed on the two large cell groups (non‐cleaned in slices, n = 19 and acutely dissociated, n = 19), however, showed that the time constant of recovery from high [Cl^−^]_i_ was neither significantly correlated to the membrane capacitance nor to leak conductance or resting potential when studied within each preparation (i. e. Spearman′s rank correlation coefficient was not significantly different from zero.) Thus, differences in resting membrane properties *per se* are unlikely to account for the differences in time course of [Cl^−^]_i_ recovery. The lack of correlation between [Cl^−^]_i_ recovery and leak conductance is in agreement with the experimental design where contribution of passive Cl^−^ flux is minimized by keeping the membrane voltage close to *E*_Cl_ .

A summary of ChABC effects on neuronal properties is shown in Table 1. A correlation analysis showed that that neither resting potential, leak conductance, membrane capacitance, resting [Cl^−^]_I_, nor [Cl^−^]_I_ recovery time constant were significantly correlated with the series resistance. (As expected, leak conductance was strongly positively correlated with membrane capacitance: Spearman’s rank correlation coefficient 0.681, p = 0.0013.)

**Table 1.** Summary of main resting neuronal properties, [Cl^−^]_i_ recovery time constant and the effect of ChABC treatment on those properties. The far right column gives p‐values from a Mann‐Whitney test for difference between control conditions and ChABC treatment. Note that only resting [Cl^−^]_i_ and [Cl^−^]_i_ recovery time constant were significantly affected. The control values reported were obtained from cells pre‐incubated at at ~ 36 °C for 2 h to enable comparison with cells from ChABC‐treated slices. N.S., not significant; *U*_rest_, resting potential; *g*_leak_, leak conductance; *C*_m_, membrane capacitance; [Cl^−^]_i_rest_, resting [Cl^−^]_I_ ; *τ*_recovery_, [Cl^−^]_I_ recovery time constant.

|  | Control | n | ChABC | n | p / N.S. |
| --- | --- | --- | --- | --- | --- |
| $U_{\text{rest}}$ | $-70 \pm 2$ mV | 19 | $-67 \pm 2$ mV | 20 | N.S. |
| $g_{\text{leak}}$ | $1.5 \pm 0.1$ nS | 19 | $1.3 \pm 0.2$ nS | 18 | N.S. |
| $C_m$ | $17.6 \pm 1.8$ pF | 21 | $15.1 \pm 2.2$ pF | 19 | N.S. |
| $[\text{Cl}^-]_{i_{\text{rest}}}$ | $6.6 \pm 0.4$ mM | 18 | $10.0 \pm 1.1$ mM | 17 | $1.7 \cdot 10^{-3}$ |
| $\tau_{\text{recovery}}$ | $137 \pm 18$ s | 19 | $441 \pm 84$ s | 15 | $4.8 \cdot 10^{-6}$ |

### ECM degradation is accompanied by calpain‐mediated reduction of KCC2 protein

Next, we used Western blot analysis to test whether the dependence of KCC2 function and of [Cl^−^]_i_ on the ECM is mediated by an altered amount of KCC2 protein. Indeed, ChABC significantly reduced the total level of KCC2 protein (to 73 ± 8 %; n = 13, p = 0.0081) but did not significantly affect the level of transferrin receptor (p = 0.57, n = 12), another membrane protein (Fig. 5*A*). Since KCC2 function may depend on the amount of di‐ or oligomers in the membrane (Blaesse et al., 2006; Watanabe et al., 2009), we also tested the hypothesis that ChABC differentially affects KCC2 monomers and dimers. ChABC significantly reduced the amount of KCC2 monomers (to 56 ± 7 %; n = 12, p = 9.8 10^−4^) but the slight apparent reduction (to 84 ± 10 %; n = 12) of dimers was not significant (p = 0.13; Fig. 5*B*), suggesting that the monomeric form is of functional importance for Cl^−^ transport sensitive to ChABC.

**Figure 5.**
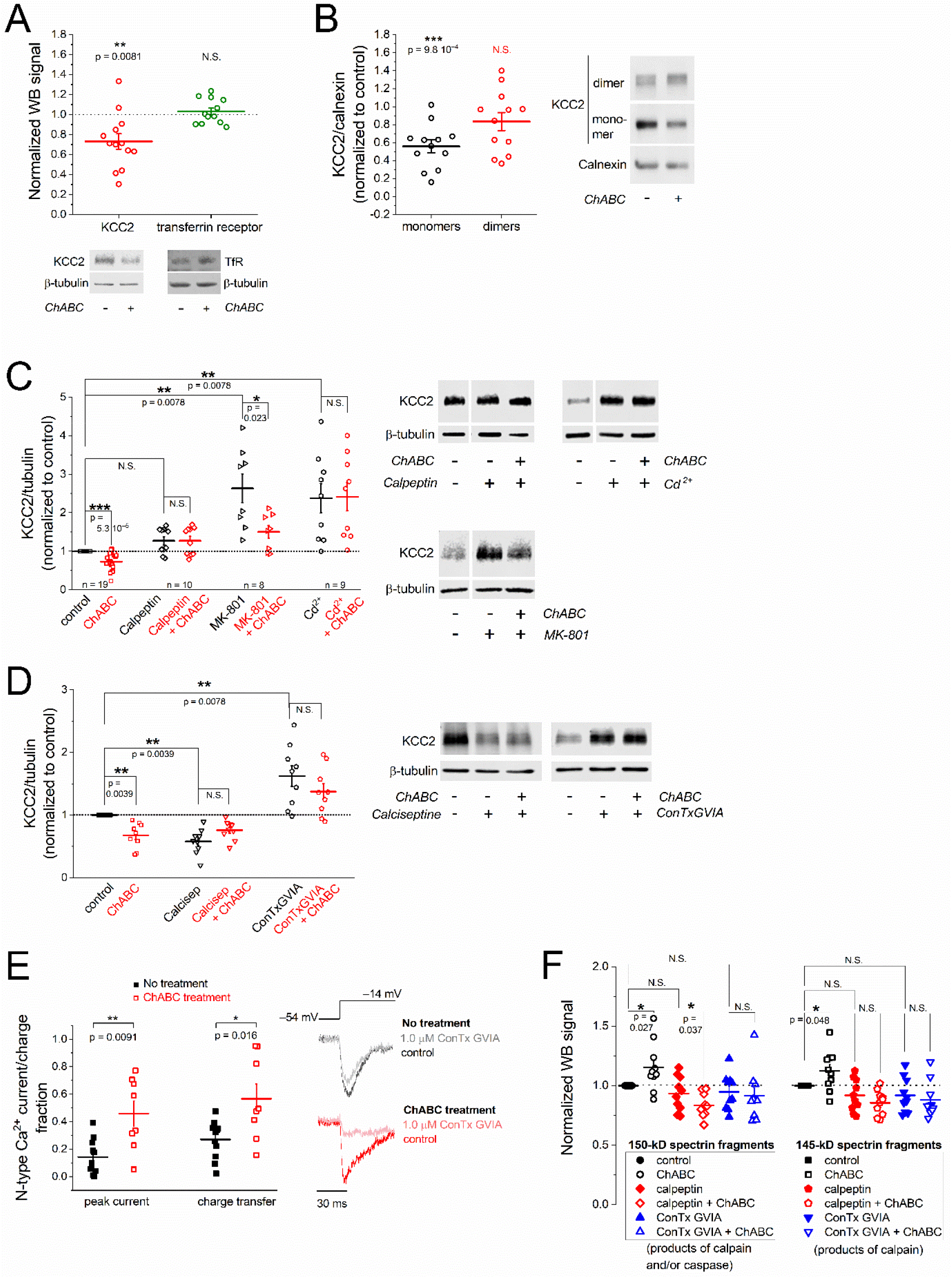
KCC2 protein level depends on the ECM, calpain and Ca^2+^ channels. *(Legend to Fig. 5, continued)* ***A***, Top: Effect of ChABC treatment on KCC2 and transferrin‐receptor (TfR) protein levels relative to β‐tubulin (β‐ tub) in whole‐cell lysate. Data normalized to control. (WB: Western blot.) Bottom: representative WB. ***B***, Left: Effect of ChABC treatment on KCC2 monomer and dimer levels in membrane fraction, relative to calnexin. Right: representative WB. ***C*** – ***D***, Left: Effects of calpeptin (30 µM), MK‐801 (1.0 µM) and Cd^2+^ (200 µM) (***C***) and of calciseptine (Calcisep; 500 nM) and ω‐conotoxin GVIA (ConTxGVIA; 1.0 µM) (***D***) on the KCC2 protein level in whole‐cell lysate in the presence or absence of ChABC, estimated as in ***A***. Right: representative WB. ***E***, ChABC effects on N‐type Ca^2+^ channels. Current contributed by N‐type Ca^2+^ channels, estimated as ω‐conotoxin GVIA‐sensitive fraction of peak Ca^2+^ current (left two symbol sets) and of charge transfer (right two symbol sets), during the first 100 ms after command voltage steps from –54 mV to –14 mV. Control conditions (filled symbols) and after ChABC treatment (similar as for protein analysis; open symbols). Example raw current traces during voltage step (top) shown in the right panel. To enable comparison between cells with different Ca^2+^ current magnitudes, only relative, ω‐conotoxin GVIA‐sensitive, current and charge fractions were used in the analysis of ChABC effects. Note that significantly larger fractions of current and charge transfer are sensitive to ω‐conotoxin GVIA after ChABC treatment. ***F***, ChABC effects on αII‐spectrin degradation. Normalized (to control) WB signals for 150 kD (left) and 145 kD (right) fragments of αII‐spectrin in whole‐cell lysate in control conditions and after treatment with ChABC, calpeptin, ω‐conotoxin GVIA (ConTx GVIA) and combinations of ChABC and calpeptin/ConTx GVIA as indicated. Levels of αII‐spectrin are relative to β‐tubulin. Bars indicate mean ± S.E.M. Statistical significance was evaluated with the paired Wilcoxon signed rank test.

KCC2 may be cleaved by calpain, a Ca^2+^‐dependent protease which has been linked to synaptic plasticity and neurodegeneration (Baudry et al., 2013), likely as a consequence of Ca^2+^ influx through NMDA receptors (Chamma et al., 2013; Puskarjov et al., 2012). Therefore, we tested the hypothesis that the effect of ECM removal on KCC2 protein level is mediated by calpain activity. In the presence of the calpain‐inhibitor calpeptin (30 µM; Sigma‐Aldrich), ChABC failed to reduce the KCC2 level (Fig. 5*C*; p = 1.0), suggesting that ECM removal triggers calpain‐dependent cleavage of KCC2.

### KCC2 downregulation is mediated by Ca^2+^ influx through N‐type channels

The KCC2 level was dramatically increased (to 263 ± 37 %; n = 8, p = 0.0078) in the presence of the NMDA‐receptor blocker MK‐801 (1.0 µM; Sigma‐Aldrich) when compared with control (Fig. 5*C*), implying a role for NMDA receptors in maintaining the basal KCC2 level. However, ChABC significantly reduced the KCC2 level (by 35 ± 10 %; n = 8; p = 0.023) also in the presence of MK‐801, contradicting the hypothesis that ECM degradation affects KCC2 via Ca^2+^ influx through NMDA receptors.

Another possible source of Ca^2+^ influx, that hypothetically may trigger calpain activation upon ECM degradation, is voltage‐gated Ca^2+^ channels. This idea was supported by the findings that in the presence of the high‐threshold Ca^2+^‐channel blocker Cd^2+^ (200 µM) (Sundgren‐Andersson and Johansson, 1998), the level of KCC2 was markedly increased (to 238 ± 38 %; n = 9, p = 0.0078) compared with control, and not significantly (p = 0.65, n = 9) affected by ChABC (Fig. 5*C*). To identify the involved channel types, the effect of ChABC treatment on the level of KCC2 was studied in the presence of calciseptine (500 nM; Latoxan) and ω‐conotoxin GVIA (1.0 µM; Alomone labs), selective blockers of L‐ and N‐type voltage‐gated Ca^2+^ channels, respectively. Surprisingly, incubation in calciseptine *per se* significantly *reduced* KCC2 (by 42 ± 7 %; n = 9, p = 0.0039), to a level similar to that in ChABC alone (p = 0.57) and also similar to the KCC2 level in a combination of ChABC and calciseptine (p = 0.098; Fig. 5*D*). The effect of calciseptine suggests that Ca^2+^ entry via L‐type channels is not coupled to the KCC2 reduction via calpain but on the contrary contributes to KCC2 upregulation. ω‐conotoxin GVIA, however, induced an opposite effect, significantly increasing the KCC2 level (to 162 ± 16 %; n = 9, p = 0.0078) when incubated without ChABC. Importantly, when co‐ applied with ω‐conotoxin GVIA, ChABC did not significantly reduce the KCC2 level below that in ω‐ conotoxin GVIA alone (p = 0.50; Fig. 5*D*). This suggests that Ca^2+^ influx through N‐type channels does contribute to the KCC2 downregulation upon ECM degradation.

Whole‐cell Ca^2+^ currents recorded under voltage‐clamp conditions (see Materials and Methods) confirmed that the ω‐conotoxin GVIA‐sensitive, presumably N‐type channel‐mediated, fraction of high‐voltage activated Ca^2+^ current was significantly larger after treatment with ChABC (Fig. 5*E*). Thus, the ω‐conotoxin GVIA‐sensitive fraction contributed 14 ± 4 % of the peak current in control conditions but increased to 46 ± 9 % after ChABC treatment (p = 0.0091, control: n = 11, ChABC: n = 8, Mann‐Whitney test) and the corresponding charge transfer (during 100 ms) increased from 27 ± 4 % to 57 ± 11 % (p = 0.016, Mann‐Whitney test).

The results above suggest that the ChABC‐induced reduction of KCC2 protein level is mediated via calpain as a consequence of Ca^2+^ influx through N‐type channels. To confirm that ChABC treatment activates calpain, we analyzed the effect of ChABC on αII‐spectrin degradation, commonly used as indicator of calpain activation (Wang, 2000), by Western blot. Spectrin fragments of 150 kDa, products of calpain or caspase‐3 activity, 145 kDa, products of calpain activity, and 120 kDa, products of caspase‐3 activity (Nath et al., 1996), were detected in control tissue and were neither significantly affected by tissue exposure to calpeptin (n = 10) nor by exposure to ω‐conotoxin GVIA (n = 10). After treatment with ChABC, the levels of 150 and 145 kDa fragments, but not that of 120 kDa fragments, were significantly increased (n = 10, Fig. 5*F*), as expected from ChABC‐induced calpain activation. This increase was blocked by calpeptin (n = 10, Fig. 5*F*), confirming that it was caused by calpain activity. The ChABC‐induced increase of 150 and 145 kDa spectrin fragments was also blocked by ω‐conotoxin GVIA (n = 10, Fig. 5*F*), confirming the requirement of N‐type Ca^2+^ channels for the activation of calpain.

### ECM degradation may turn GABA effect from inhibition into excitation

Finally, we tested the hypothesis that the effects of ECM degradation on KCC2 function and on [Cl^−^]_i_ are relevant for GABA_A_‐receptor mediated transmission. In control slices, presynaptic stimulation was followed by inhibition of spontaneously firing cells (Fig. 6*A*, top trace). This inhibition was abolished by the GABA_A_‐receptor antagonist gabazine (SR‐95531, Tocris; n = 8, Fig. 6*A*, lower trace). In ChABC‐treated slices, spontaneous firing frequency was, neither before nor after (0 – 40 ms) stimulation, significantly different from control. A sequence of high‐frequency stimulation (HFS; 500 stimuli at 40 Hz), however, significantly reduced the inhibitory effect of 0.4‐Hz stimulation in ChABC‐treated cells (n = 22, p = 0.0061, paired Wilcoxon signed ranks test), but not in control cells (n = 13, p = 0.20, paired Wilcoxon signed rank test; Fig. 6*B*). Ten minutes after HFS, the change was accentuated and an initial excitatory effect (Fig. 6*C*) was seen in several ChABC‐exposed cells upon 0.4‐Hz stimulation, resulting in average excitation in ChABC‐exposed cells (Fig. 6*D*), but average inhibition in control cells (Fig. 6*D*) 0 – 40 ms after stimulation. The reduction in immediate post‐HFS synaptic inhibition of ChABC‐exposed cells (Fig. 6*B*, middle) was abolished (p = 0.45, n = 18, paired Wilcoxon signed ranks test) when the calpain blocker calpeptin (30 μM) was present during slice incubation with ChABC (Fig. 6*B*, right), and synaptic inhibition 10 minutes after HFS was rescued (Fig. 6*D*, right; p = 0.27, n = 18, paired Wilcoxon signed ranks test). These effects of calpeptin on synaptic transmission are consistent with our biochemical analysis of KCC2 and calpain activity and imply that calpain transduces attenuation of ECM into degradation of KCC2 and thereby increases [Cl^−^]_i_ and modifies GABA‐mediated transmission.

**Figure 6.**
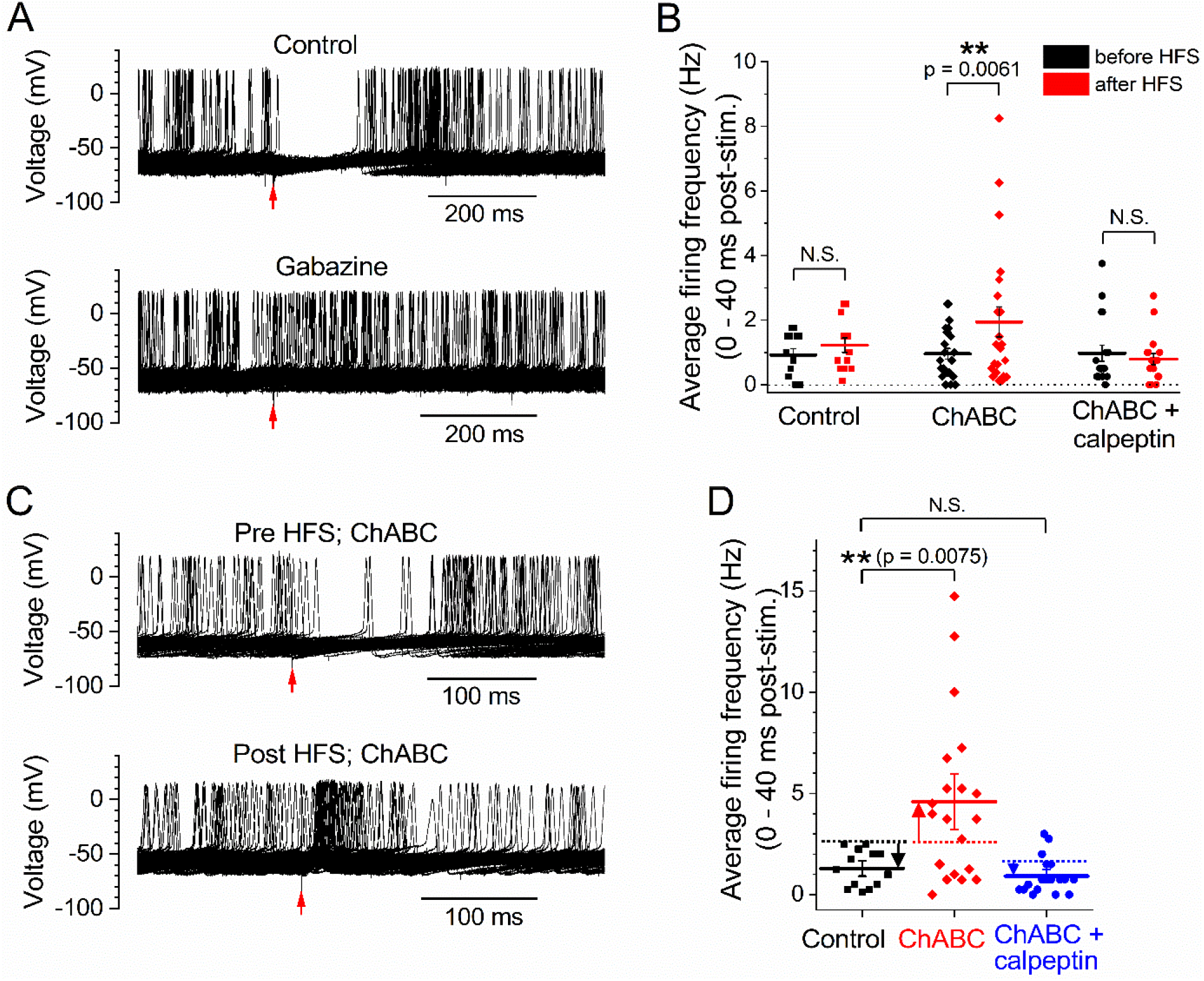
Synaptic transmission after high‐frequency stimulation is altered by ECM degradation. ***A***, Presynaptic stimulation (arrow) at 0.4 Hz in the presence of glutamate‐receptor blockers (20 μM NBQX + 50 μM APV) inhibits spontaneous firing (top; 100 superimposed traces). After addition of the GABA_A_‐receptor antagonist gabazine (SR‐95531; 3.0 μM), similar stimulation does not inhibit spontaneous firing (bottom; 100 superimposed traces). ***B***, A period of high‐frequency stimulation (HFS; 500 stimuli at 40 Hz) does not change the inhibitory effect of 0.4‐Hz stimulation on firing frequency in cells from control slices (left) but reduces the inhibitory effect in cells from ChABC‐treated slices (middle). This effect of ChABC is abolished by the calpain inhibitor calpeptin (30 μM; right). (Horizontal lines and error bars: mean ± S.E.M.) ***C***, Top: Hundred superimposed traces showing initial inhibitory effect of 0.4‐Hz stimulation (arrow) on a cell in a ChABC‐treated slice. Bottom: Excitatory effect of similar stimulation in the same cell but 10 minutes after HFS (500 stimuli at 40 Hz). ***D***, Ten minutes after HFS, the change in transmission is accentuated in many ChABC‐exposed cells, and frequently the initial inhibition (left) has changed to excitation (middle). Calpeptin (30 μM) prevents the loss of inhibition in ChABC‐treated slices (right). Data show impulse frequency 0 – 40 ms after stimulus (repeated at 0.4 Hz). Solid horizontal lines and error bars: mean ± S.E.M. Dotted lines show pre‐stimulation mean frequency; arrows indicate the change to mean post‐stimulus (0 – 40 ms) frequency, downward for inhibition (control and ChABC + calpeptin) and upward for excitation (ChABC).

## Discussion

In conclusion, the present study verifies the recently suggested importance of the ECM for neuronal Cl^−^ homeostasis (Glykys et al., 2014a) and uncovers a novel molecular mechanism for control of GABA‐mediated transmission. Direct electrophysiological measurement showed that basal [Cl^−^]_i_ is lower than expected for a passive distribution and depends on KCC2 and on ECM integrity. However, the results are not consistent with the debated mechanism involving a strong effect of ECM negative charges on [Cl^−^]_o_ and [Cl^−^]_i_ (Glykys et al., 2014a). Thus, ECM degradation does not affect [Cl^−^]_o_, but affects KCC2, mainly in monomeric form, and basal [Cl^−^]_i_. Importantly, the rate of KCC2‐mediated Cl^−^ extrusion after a high load, critical to neural coding (Doyon et al., 2016), also depends on ECM integrity. The mechanism by which the ECM degradation influences KCC2 function, largely depending on the Ca^2+^‐activated protease calpain, partly overlaps with that for the glutamate‐evoked reduction in KCC2 associated with interictal‐like activity (Chamma et al., 2013; Puskarjov et al., 2012). However, our results do not support a role of NMDA‐receptors in the effect of ECM removal. Rather, they suggest that ECM degradation reduces KCC2 level via Ca^2+^ entry through N‐type Ca^2+^ channels and subsequent calpain activation. A role for calpain in increasing excitability via reduced KCC2 and increased [Cl^−^]_i_ is particularly intriguing since calpain is activated at brain injury and during epileptogenesis (Lam et al., 2017). The present findings therefore suggest a positive feedback mechanism by which epileptogenesis may be self‐sustaining, with seizure‐induced calpain activation contributing to further increased cell excitability.

Our findings also indicate a novel differential and opposing regulation of KCC2 via different Ca^2+^ channel subtypes. The suggested upregulation of KCC2 level via Ca^2+^ entry through L‐type channels is consistent with the roles of L‐type channels and Ca^2+^/calmodulin‐dependent protein kinase II for elevation of KCC2 transcription (Li et al., 2016) and of KCC2 protein level (Liedtke et al., 2013). Since removal of the major ECM component hyaluronic acid reduces Ca^2+^ influx through L‐type channels in hippocampal CA1 cells (Kochlamazashvili et al., 2010), this may provide a calpain‐independent mechanism by which the ECM regulates KCC2. In principle, the N‐type channel‐ and calpain‐dependent pathway and the L‐type channel‐dependent, calpain‐independent pathway may act in concert to reduce the KCC2 level upon ECM degradation, but apparently the former pathway dominates the effect of ChABC studied here.

Finally, the present results show that the ECM is critically important for synaptic function, since ECM degradation may change a dominant inhibitory effect to excitatory. Similar changes in excitability have been described for other conditions with intracellular Cl^−^ accumulation (Cohen et al., 2002; Coull et al., 2003; Khazipov et al., 2015). We speculate that in conditions, such as epilepsy, where the ECM may be downregulated (Soleman et al., 2013), effects on [Cl^−^]_i_ may contribute to the pathology. Furthermore, enzymatic degradation of ECM has been shown to re‐open a “window” of enhanced developmental plasticity and aided functional recovery in regeneration, stroke and amblyopia (Soleman et al., 2013) and calpain inhibition ameliorates seizure burden in an epilepsy model (Lam et al., 2017). Further studies are warranted to explore the possibility that these therapeutic effects at least partially are mediated by [Cl^−^]_i_‐dependent changes in GABAergic transmission.

## Conclusions

In this work, we show that degradation of the extracellular matrix (ECM) which surrounds brain cells increases basal [Cl^‐^]_i_, and reduces the rate of Cl^‐^ extrusion via a reduced amount of the Cl^‐^ transporting protein K‐Cl‐cotransporter 2 (KCC2). This results in a reduced inhibitory, or even an excitatory, effect of GABA‐mediated transmission. The findings imply a novel pathway for the control of neuronal [Cl^‐^]_i_ and excitability and may explain how degradation of ECM may re‐open a window for developmental forms of neuroplasticity in the adult central nervous system.

## Acknowledgements

This work was supported by the Swedish Research Council (grant no 22292), DFG grant Po 732, Excellence Cluster REBIRTH, DFG Project‐ID 425899996 – SFB 1436 (TP05), Gunvor och Josef Anérs Stiftelse and by Umeå University Medical Faculty (Insamlingsstiftelsen and Leila och Bertil Ehrengrens fond). VU0255011‐1 was kindly provided as a gift by Dr. Craig Lindsley.

## Notes

Conflict of Interest: The authors declare no competing financial interests.

### Competing Interest Statement

The authors have declared no competing interest.

## References

Abe Y, Furukawa K, Itoyama Y, Akaike N (1994) Glycine response in acutely dissociated ventromedial hypothalamic neuron of the rat: new approach with gramicidin perforated patch‐clamp technique. J Neurophysiol 72:1530–1537.

Baudry M, Chou MM, Bi X (2013) Targeting calpain in synaptic plasticity. Expert Opin Ther Targets 17:579–592.

Ben‐Ari Y (2014) The GABA excitatory/inhibitory developmental sequence: a personal journey. Neuroscience 279:187–219. doi:10.1016/j.neuroscience.2014.08.001

Ben‐Ari Y (2002) Excitatory actions of GABA during development: the nature of the nurture. Nat Rev Neurosci 3:728–739.

Blaesse P, Guillemin I, Schindler J, Schweizer M, Delpire E, Khiroug L, Friauf E, Nothwang HG (2006) Oligomerization of KCC2 correlates with development of inhibitory neurotransmission. J Neurosci 26:10407–10419.

Bormann J, Hamill OP, Sakmann B (1987) Mechanism of anion permeation through channels gated by glycine and gamma‐aminobutyric acid in mouse cultured spinal neurones. J Physiol 385:243–286.

Boron WF (2009a) Organization of the respiratory system. In: Medical physiology (Boron WF, Boulpaep EL, eds), pp613–629. Philadelphia: Saunders, Elsevier.

Boron WF (2009b) Acid‐base physiology. In: Medical physiology (Boron WF, Boulpaep EL, eds), pp652–671. Philadelphia: Saunders, Elsevier.

Bukalo O, Schachner M, Dityatev A (2001) Modification of extracellular matrix by enzymatic removal of chondroitin sulfate and by lack of tenascin‐R differentially affects several forms of synaptic plasticity in the hippocampus. Neuroscience 104:359–369.

Butler JN (1964) Ionic equilibration. A mathematical approach. Reading, Massachusetts: Addison‐ Wesley.

Chamma I, Heubl M, Chevy Q, Renner M, Moutkine I, Eugène E, Poncer JC, Lévi S (2013) Activity‐ dependent regulation of the K/Cl transporter KCC2 membrane diffusion, clustering, and function in hippocampal neurons. J Neurosci 33:15488–15503.

Chavas J, Marty A (2003) Coexistence of excitatory and inhibitory GABA synapses in the cerebellar interneuron network. J Neurosci 23:2019–2031.

Chen H‐X, Otmakhov N, Lisman J (1999) Requirements for LTP Induction by pairing in hippocampal CA1 pyramidal cells. J Neurophysiol 82:526–532.

Cho C‐H (2014) Star players sidelined in chloride homeostasis in neurons. Front Cell Neurosci 8:114. doi:10.3389/fncel.2014.00114

Choi HJ, Lee CJ, Schroeder A, Kim YS, Jung SH, Kim JS, Kim DY, Son EJ, Han HC, Hong SK, Colwell CS, Kim YI (2008) Excitatory actions of GABA in the suprachiasmatic nucleus. J Neurosci 28:5450–5459.

Cohen I, Navarro V, Clemenceau S, Baulac M, Miles R (2002) On the origin of interictal activity in human temporal lobe epilepsy in vitro. Science 298:1418–1421.

Coull JAM, Boudreau D, Bachand K, Prescott SA, Nault F, Sík A, De Koninck P, De Koninck Y (2003) Trans‐synaptic shift in anion gradient in spinal lamina I neurons as a mechanism of neuropathic pain. Nature 424:938–942.

Davies CW (1962) Ion Association. London: Butterworths.

DeFazio RA, Heger S, Ojeda SR, Moenter SM (2002) Activation of A‐type gamma‐aminobutyric acid receptors excites gonadotropin‐releasing hormone neurons. Mol Endocrinol 16:2872–2891.

Delpire E, Staley KJ (2014) Novel determinants of the neuronal Cl– concentration. J Physiol 592:4099–4114.

Doyon N, Prescott SA, De Koninck Y (2016) Mild KCC2 hypofunction causes inconspicuous chloride dysregulation that degrades neural coding. Front Cell Neurosci 9. doi:10.3389/fncel.2015.00516

Edwards FA, Konnerth A, Sakmann B, Takahashi T (1989) A thin slice preparation for patch clamp recordings from neurones of the mammalian central nervous system. Pflügers Arch 414:600–612.

Freedman JC, Hoffman JF (1979) Ionic and osmotic equilibria of human red blood cells treated with nystatin. J Gen Physiol 74:157–185.

Friedman MH (2010) Principles and models of biological transport, 2nd ed. New York: Springer.

Glykys J, Dzhala V, Egawa K, Balena T, Saponjian Y, Kuchibhotla KV, Bacskai BJ, Kahle KT, Zeuthen T, Staley KJ (2014a) Local impermeant anions establish the neuronal chloride concentration. Science 343:670–675.

Glykys J, Dzhala V, Egawa K, Balena T, Saponjian Y, Kuchibhotla KV, Bacskai BJ, Kahle KT, Zeuthen T, Staley KJ (2014b) Response to comments on “Local impermeant anions establish the neuronal chloride concentration”. Science 345:1130.

Haam J, Popescu IR, Morton LA, Halmos KC, Teruyama R, Ueta Y, Tasker JG (2012) GABA is excitatory in adult vasopressinergic neuroendocrine cells. J Neurosci 32:572–582.

Härtig W, Brauer K, Brückner G (1992) Wisteria floribunda agglutinin‐labelled nets surround parvalbumin‐containing neurons. Neuroreport 3:869–872.

Howell MD, Bailey LA, Cozart MA, Gannon BM, Gottschall PE (2015) Hippocampal administration of chondroitinase ABC increases plaque‐adjacent synaptic marker and diminishes amyloid burden in aged APPswe/PS1dE9 mice. Acta Neuropathol Commun 3:54. doi:10.1186/s40478‐015‐0233‐z

Huberfeld G, Wittner L, Clemenceau S, Baulac M, Kaila K, Miles R, Rivera C (2007) Perturbed chloride homeostasis and GABAergic signaling in human temporal lobe epilepsy. J Neurosci 27:9866–9873.

Hughes DI, Bannister AP, Pawelzik H, Thomson AM (2000) Double immunofluorescence, peroxidase labelling and ultrastructural analysis of interneurones following prolonged electrophysiological recordings in vitro. J Neurosci Methods 101:107–116.

Izu YC, Sachs F (1991) Inhibiting synthesis of extracellular matrix improves patch clamp seal formation. Pflügers Arch 419:218–220.

Johansson S, Yelhekar TD, Druzin M (2016) Commentary: Chloride regulation: a dynamic equilibrium crucial for synaptic inhibition. Front Cell Neurosci 10. doi:10.3389/fncel.2016.00182

Kaila K, Price TJ, Payne JA, Puskarjov M, Voipio J (2014) Cation‐chloride cotransporters in neuronal development, plasticity and disease. Nat Rev Neurosci 15:637–654.

Kajino K, Matsumura Y, Fujimoto M (1982) [Determination of dissociation exponent of CO2 used in Henderson‐Hasselbalch equation by means of bicarbonate‐selective microelectrode]. Nihon Seirigaku Zasshi. 44:663–673.

Karlsson U, Druzin M, Johansson S (2011) Cl– concentration changes and desensitization of GABAA and glycine receptors. J Gen Physiol 138:609–626. doi:10.1085/jgp.201110674

Khazipov R, Valeeva G, Khalilov I (2015) Depolarizing GABA and developmental epilepsies. CNS Neurosci Ther 21:83–91.

Kochlamazashvili G, Henneberger C, Bukalo O, Dvoretskova E, Senkov O, Lievens PM‐J, Westenbroek R, Engel AK, Catterall WA, Rusakov DA, Schachner M, Dityatev A (2010) The extracellular matrix molecule hyaluronic acid regulates hippocampal synaptic plasticity by modulating postsynaptic L‐type Ca2+ channels. Neuron 67:116–128.

Kyrozis A, Reichling DB (1995) Perforated‐patch recording with gramicidin avoids artifactual changes in intracellular chloride concentration. J Neurosci Methods 57:27–35.

Lam PM, Carlsen J, González MI (2017) A calpain inhibitor ameliorates seizure burden in an experimental model of temporal lobe epilepsy. Neurobiol Dis 102:1–10.

Li B, Tadross MR, Tsien RW (2016) Sequential ionic and conformational signaling by calcium channels drives neuronal gene expression. Science 351:863–867.

Liedtke W, Yeo M, Zhang H, Wang Y, Gignac M, Miller S, Berglund K, Liu J (2013) Highly conductive carbon nanotube matrix accelerates developmental chloride extrusion in central nervous system neurons by increased expression of chloride transporter KCC2. Small 9:1066–1075.

Luhmann HJ, Kirischuk S, Kilb W (2014) Comment on “Local impermeant anions establish the neuronal chloride concentration”. Science 345:1130.

Mahadevan V, Pressey JC, Acton BA, Uvarov P, Huang MY, Chevrier J, Puchalski A, Li CM, Ivakine EA, Airaksinen MS, Delpire E, McInnes RR, Woodin MA (2014) Kainate receptors coexist in a functional complex with KCC2 and regulate chloride homeostasis in hippocampal neurons. Cell Rep 7:1762–1770.

Malinina E, Druzin M, Johansson S (2005) Fast neurotransmission in the rat medial preoptic nucleus. Brain Res 1040:157–168.

Morita S, Oohira A, Miyata S (2010) Activity‐dependent remodeling of chondroitin sulfate proteoglycans extracellular matrix in the hypothalamo‐neurohypophysial system. Neuroscience 166:1068–1082.

Moyer JR, Brown TH (2002) Patch‐clamp techniques applied to brain slices. In: Neuromethods 35: Patch‐clamp analysis: Advanced techniques (Walz W, Boulton AA, Baker GB, eds), pp 135–193. Totowa, New Jersey: Humana.

Myers VB, Haydon DA (1972) Ion transfer across lipid membranes in the presence of gramicidin A. II. The ion selectivity. Biochim Biophys Acta 274:313–322.

Nath R, Raser KJ, Stafford D, Hajimohammadreza I, Posner A, Allen H, Talanian RV, Yuen P, Gilbertsen RB, Wang KKW (1996) Non‐erythroid α‐spectrin breakdown by calpain and interleukin 1β‐ converting‐enzyme‐like protease(s) in apoptoti cells: contributory roles of both protease families in neuronal apoptosis. Biochem J 319:683–690.

Pusch M, Neher E (1988) Rates of diffusional exchange between small cells and a measuring patch pipette. Pflügers Arch 411:204–211.

Puskarjov M, Ahmad F, Kaila K, Blaesse P (2012) Activity‐dependent cleavage of the K‐Cl cotransporter KCC2 mediated by calcium‐activated protease calpain. J Neurosci 32:11356–11364.

Savtchenko LP, Poo MM, Rusakov DM (2017) Electrodiffusion phenomena in neuroscience: a neglected companion. Nat Rev Neurosci 18:598–612.

Senkov O, Andjus P, Radenovic L, Soriano E, Dityatev A (2014) Neural ECM molecules in synaptic plasticity, learning, and memory. Prog Brain Res 214:53–80.

Soleman S, Filippov MA, Dityatev A, Fawcett JW (2013) Targeting the neural extracellular matrix in neurological disorders. Neuroscience 253:194–213.

Stumm W, Morgan JJ (1996) Aquatic chemistry: Chemical equilibria and rates in natural waters. 3rd ed. New York: John Wiley & Sons.

Sundgren‐Andersson AK, Johansson S (1998) Calcium spikes and calcium currents in neurons from the medial preoptic nucleus of rat. Brain Res 783:194–209.

Voipio J, Boron WF, Jones SW, Hopfer U, Payne JA, Kaila K (2014) Comment on “Local impermeant anions establish the neuronal chloride concentration”. Science 345:1130.

Wang KKW (2000) Calpain and caspase: can you tell the difference? Trends Neurosci 23:20–26.

Watanabe M, Wake H, Moorhouse AJ, Nabekura J (2009) Clustering of neuronal K+‐Cl– cotransporters in lipid rafts by tyrosine phosphorylation. J Biol Chem 284:27980–27988.

Williams JR, Payne JA (2004) Cation transport by the neuronal K+‐Cl– cotransporter KCC2: thermodynamics and kinetics of alternate transport modes. Am J Physiol Cell Physiol 287:C919–C931.

Yelhekar TD, Druzin M, Karlsson U, Blomqvist E, Johansson S (2016) How to properly measure a current‐voltage relation?—Interpolation vs. ramp methods applied to Studies of GABAA receptors. Front Cell Neurosci 10. doi:10.3389/fncel.2016.00010

Yelhekar TD, Druzin M, Johansson S (2017) Contribution of resting conductance, GABAA‐receptor mediated miniature synaptic currents and neurosteroid to chloride homeostasis in central neurons. eNeuro 4:e0019–17.2017. doi: 10.1523/ENEURO.0019–17.2017

